# Conformational Expansion Underlies the Evolutionary Emergence of Redox Sensitivity in Vertebrate Glucokinases

**DOI:** 10.64898/2026.05.13.724885

**Authors:** Johanna E. Papa, A. Carl Whittington, Brian G. Miller

## Abstract

Glucokinase (GCK) catalyzes the first step of glycolysis in pancreatic β-cells, where it functions as the body’s primary glucose sensor. GCK is extremely sensitive to oxidative inactivation, both *in vivo* and *in vitro*. This characteristic provides a mechanism to regulate GCK activity via alterations in the cellular redox environment. To understand the molecular and evolutionary origins of redox regulation, we characterized the sensitivity to oxidative inactivation of four extant GCKs and five ancestral GCKs produced from a recent phylogenetic analysis of the vertebrate family. We find that two invertebrate GCKs are significantly less sensitive to oxidative inactivation compared to their vertebrate counterparts. We also demonstrate that an ancestral GCK from chordates (cGCK) is insensitive to oxidative inactivation, whereas an ancestral GCK from early vertebrates (vGCK) displays a degree of redox responsiveness comparable to the extant human enzyme. The redox insensitive cGCK ancestor lacks cysteine residues at two positions, Cys230 and Cys461, that are conserved in all redox sensitive ancestral and extant enzymes. We find that installation of cysteines at these positions is insufficient to install redox sensitivity into cGCK. Our data demonstrate that the appearance of redox responsiveness in GCKs coincides with an expansion in the conformational landscape of the protein that occurred during early vertebrate evolution. These observations support a model in which the emergence of redox sensitivity required the ability to sample a unique super-open conformation, an event that also facilitated the emergence of two orthogonal GCK regulatory strategies, allosteric regulation by substrate glucose and an inhibitory interaction with the glucokinase regulatory protein.

## INTRODUCTION

Glucokinase (GCK) plays an important role in the human body by sensing and responding to alterations in blood glucose concentrations. As such, this enzyme is subject to a variety of functionally distinct regulatory mechanisms that govern its cellular activity.^1^ GCK displays a cooperative kinetic response to substrate glucose, with a midpoint value that approximates physiological blood glucose levels.^2,3^ This unique form of substrate-mediated autoregulation, reflected in a hill coefficient of 1.7, originates from an ability of the unliganded enzyme to sample multiple conformations.^4,5^ Indeed, kinetic cooperativity and conformational heterogeneity are tightly linked, and the magnitude of the hill coefficient is a proxy for conformational diversity among GCKs. In the liver, GCK is additionally regulated via an inhibitory protein-protein interaction with the glucokinase regulatory protein (GKRP), which serves to sequester the enzyme in the hepatocyte nucleus under conditions of low glucose.^6^ Beyond these regulatory mechanisms, posttranslational modification of GCK via ubiquitination, sumoylation and nitrosylation provides further control over the stability, localization and enzymatic activity of the protein.^1^

Mammalian GCK is highly sensitive to the reduction potential of its surrounding environment.^7–11^ In the absence of exogenous reductants and cognate ligands, GCK undergoes rapid inactivation via a process that reduces the enzyme’s apparent *k*_cat_ value, but which leaves the glucose *K*_0.5_ value unaltered.^7^ Addition of glucose alters this inactivation process to one in which the glucose *K*_0.5_ value shifts to higher values over time, while the *k*_cat_ value remains unaltered.^7^ Addition of small-molecule reductants restores wild-type kinetic characteristics and protects the enzyme against both types of inactivation.^7^ Mechanistic studies of the redox sensitivity of rat liver GCK support a model in which oxidation of one or more cysteine residues alters the distribution of protein conformations that constitute the native state ensemble.^7,8,10–14^ Protection against oxidative inactivation via the addition of glutathione, the primary thiol-based reductant in eukaryotic cells, is concentration dependent with a midpoint value of 4 mM.^7^ This value lies within the reported cellular levels of glutathione^7^, suggesting that redox regulation of glucokinase represents a physiologically relevant mechanism to control GCK activity.

The amino acid sequence of human GCK (hGCK) harbors 12 total cysteine residues that are distributed throughout the enzyme’s three-dimensional structure (Figure 1). Four of these residues, Cys220, Cys364, Cys371 and Cys434, represent consensus S-nitrosylation sites.^8^ Individual replacement of these four residues with serine identified position 371 as the primary site of S-nitrosylation of GCK *in vivo*.^8^ A subsequent investigation using Förster resonance energy transfer supported a model in which S-nitrosylation of GCK activates the enzyme by promoting a redistribution of enzyme conformations toward a state that resembles the glucose-bound state.^8^ In a separate study, partial or complete loss of enzyme activity was observed upon substituting serine for cysteine at seven distinct positions with human pancreatic GCK.^10^ This study also demonstrated that the removal of individual cysteine residues was insufficient to protect the enzyme against inhibition by the redox active small molecule alloxan, which stimulates the formation of multiple intramolecular disulfide bonds in GCK.^10^ Together, these findings are consistent with a model in which the conformational distribution and/or activity of GCK is linked to the structure and oxidation status of one or more cysteine side chains, thereby conferring redox responsiveness to this important regulatory enzyme.

**Figure 1.**
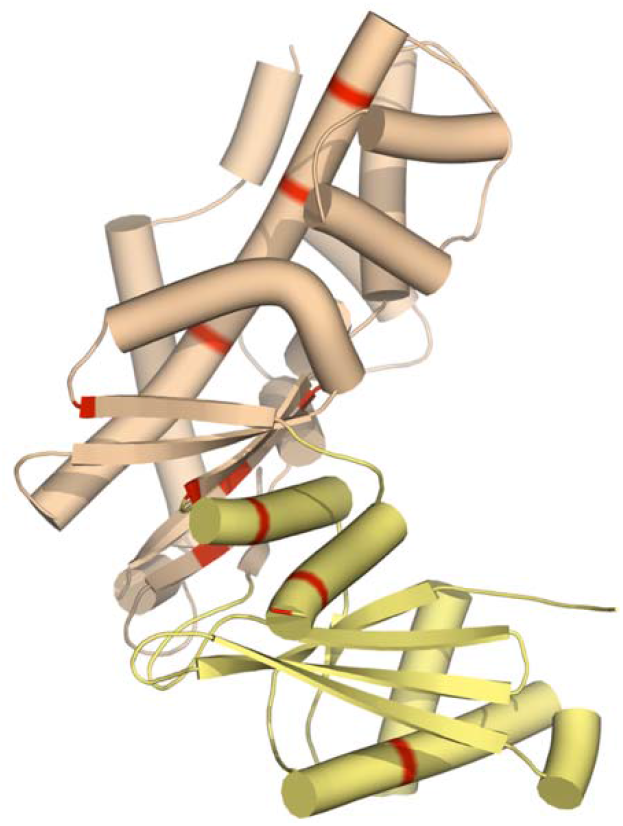
Structure of unliganded human GCK (PDB 1V4T) showing the distribution of cysteine residues (red). The large domain is tan and the small domain is yellow.

Several disease phenotypes are caused by mutations within the *gck* locus.^2^ Activating mutations that decrease cooperativity and increase glucose responsiveness are associated with persistent hyperinsulinemic hypoglycemia of infancy (PHHI). Conversely, loss-of-function inactivating substitutions cause maturity-onset diabetes of the young type II (MODY-II). Cullen and coworkers investigated the impact of oxidative stress upon both activating and inactivating disease-linked mutations using a MIN6, mouse insulinoma model system.^9^ These experiments demonstrated that MODY-associated GCK variants displayed an increased sensitivity to oxidation, whereas the redox sensitivity of PHHI-associated variants were comparable to the wild-type enzyme.^9^ MODY variants also displayed an increased sensitivity to alloxan inhibition and experienced a greater increase in activity upon exposure to dithiothreitol compared to wild-type GCK.^9^ These findings, together with cellular studies from several other investigations, support the postulate that redox regulation is an important physiological mechanism governing GCK function.

Our laboratory is interested in elucidating the molecular and evolutionary origins of the rich diversity of regulatory mechanisms found in modern GCKs. Past phylogenetic work established that GCK’s kinetic cooperativity and its inhibitory interaction with GKRP evolved concurrently along the vertebrate evolutionary trajectory, both resulting from the expansion of conformational heterogeneity in the ancestral GCK.^15^ Here, we undertook a similar vertical approach to investigate the functional transitions that led to GCK’s unique redox responsiveness. We report the sensitivity to oxidative inactivation of four extant glucokinases, the results of which suggest an intrinsic difference in redox responsiveness between enzymes from invertebrate and vertebrate organisms. We also report the sensitivity to oxidative inactivation of 5 ancestral GCKs spanning the vertebrate phylogeny. Together, the results reveal a discrete functional transition early in vertebrate evolution in which the emergence of redox responsiveness coincides with the ability of the enzyme to sample a new, super-open conformation.

## Materials and Methods

### Protein Production and Purification

The amino acid sequences of all ancestral and extant glucokinases used in this study are provided in Supporting Information (Figure S1 & S2, respectively). The maximum-likelihood models of ancestral proteins were obtained from a previously published investigation, which demonstrated that each protein was phenotypically robust to phylogenetic reconstruction.^15^ GCKs were produced as N-terminal hexahistidine-tagged polypeptides from pET22(b) plasmids harboring codon-optimized genes (Genscript). BL21(DE3) *Escherichia coli* harboring individual expression plasmids were grown at 37°C until the OD_600 nm_ reached 0.5, at which time the temperature was reduced to 20°C and IPTG was added to a final concentration of 0.4 mM. Following overnight incubation, cells were harvested via centrifugation at 5,000 xg and the resulting cell pellet was resuspended in buffer A (50 mM HEPES pH 7.6, 50 mM KCl, 40 mM imidazole, 10 mM dithiothreitol (DTT), 25% v/v glycerol). Cells were lysed three times using a French press at 4°C and 1100 psi, followed by centrifugation at 25,000 xg at 4°C for 35 minutes. Lysate was passed through a 5 mL HisTrap column (Cytiva) pre-equilibrated with buffer A. The column was washed with 50 mL buffer A and protein was eluted with 25 mL of buffer A supplemented with 0.25 M imidazole. Eluate was dialyzed in a 6-8000 MWCO bag against 1 L of 50 mM HEPES, 50 mM KCl, pH 7.6, 10 mM DTT with two buffer exchanges. Protein was then concentrated via filter centrifugation to a concentration of ∼10 mg/mL. 50 μg of vitamin B12 was added as a molecular weight standard and the mixture was injected on a Superdex 200 increase 10/300 GL column (Cytiva) at a flow rate of 0.2 mL/min. Fractions containing the highest A_280nm_ values were pooled for activity assays.

### Site-Directed Mutagenesis

Agilent’s Quikchange primer design tool was used to design primers to generate the desired mutations. The sequences of primers used in this study are provided in Table S1. Mutagenesis was conducted using the Quickchange protocol followed by sequence validation by Sanger sequencing in the DNA sequencing facility in the Department of Biological Science at Florida State University, or via whole-plasmid sequencing (Plasmidsaurus). Variant enzymes were produced and purified using the same protocol for extant and ancestral GCKs described above.

### Enzyme Activity Assays

GCK activity was quantified via an enzyme-linked spectrophotometric assay using glucose 6-phosphate dehydrogenase (G6PDH) to couple G6P production to NADP^+^ oxidation. For time course inactivation assays, a reaction mixture was prepared containing ∼20 nM GCK, 0.2 M HEPES pH 8.0, 50 mM KCl, 10 mM ATP (adjusted to pH 7.6), 11 mM MgCl_2_, 0.5 mM NADP^+^, and ± 1 mM glutathione. Samples were incubated at RT and protected from light by foil. At a given time point, an aliquot was removed to a 1 mL cuvette containing the same reaction components supplemented with 7.5 units G6PDH. A baseline was collected at 340 nm at 25 °C for 2 min, after which time the reaction was initiated via addition of 100 mM glucose. Remaining enzyme activity at various time points were collected until the final activity was reduced by 95% or until 96 hours had passed. The resulting inactivation profiles were fit to a single exponential decay equation: % activity = (% initial activity)e^-*k*t^, where *k* is the first order rate constant for inactivation. Half-lives for inactivation were calculated from rate constants using the equation t_1/2_ = ln2/*k*. An unpaired Student’s T-test was performed in GraphPad prism to determine statistical significance between half-lives for inactivation.

For investigations involving variant enzymes prepared via site-directed mutagenesis, kinetic parameters were determined by performing assays under the conditions described above except using variable ATP (0.25-10 mM) or variable glucose (0.1-100 mM). Data were fit to either the Michaelis-Menten or Hill equations depending on the enzyme under investigation. All assays were performed on two or more bioreplicates with two or more technical replicates.

## Results and Discussion

### Vertebrate GCKs display high redox sensitivity

To initiate our investigations, we sought to validate the previously reported high sensitivity of hGCK to oxidative inactivation. We also wanted to quantify the ability of the physiological reductant glutathione to protect against activity loss. We incubated highly purified recombinant hGCK in the presence of varying glutathione concentrations and measured the time-dependent loss in enzyme activity over a period of 56 hours (Figure S3). Activity measurements were performed under saturating ATP and glucose concentrations. The activity vs time curves were fitted to a single exponential decay model, the rate constant for which (*k*_dec_) was used to calculate a half-life for enzyme inactivation. In the absence of glutathione, the half-life for inactivation of hGCK was 1.7 ± 0.15 h. This value was extended to 8.3 ± 0.52 h by the inclusion of 1 mM glutathione and 36.6 ± 0.4 h by the addition of 10 mM glutathione to the reaction mixture prior to incubation (Table 1, Figure S3). Based on these results, we chose to evaluate the redox sensitivity of all enzymes investigated here in the absence and presence of 1 mM glutathione.

**Table 1.** Half-lives for oxidative inactivation of extant (top) and ancestral (bottom) glucokinases in the absence and presence of 1 mM glutathione (GSH). *indicates p-value < 0.05; *n*.*s*. indicates no significant difference.

| **Extant GCK** | **- GSH (h)** | **+ GSH (h)** | **Fold change** |
| --- | --- | --- | --- |
| *B. belcheri* | 4.6 ± 0.48 | 24.5 ± 0.78 | 5.3* |
| *C. intestinalis* | 7.4 ± 0.97 | 18.8 ± 0.72 | 2.5* |
| *D. rerio* | 1.1 ± 0.04 | 3.8 ± 0.53 | 3.5* |
| *X. laevis* | 2.6 ± 0.48 | 7.7 ± 1.7 | 3.0* |
| *H. sapiens* | 1.7 ± 0.15 | 8.3 ± 0.52 | 4.9* |
| **Ancestral GCK** | **- GSH (h)** | **+ GSH (h)** | **Fold change** |
| *cGCK* | 62.5 ± 13.0 | 59.9 ± 17.0 | *n.s.* |
| *vGCK* | 3.3 ± 0.98 | 15.9 ± 2.3 | 4.8* |
| *gGCK* | 8.3 ± 1.6 | 11.1 ± 0.32 | 1.3* |
| *tGCK* | 4.7 ± 0.81 | 12.3 ± 2.6 | 2.6* |
| *mGCK* | 1.1 ± 0.18 | 5.4 ± 2.7 | 4.9* |

Next, we examined the redox sensitivity of enzymes from taxonomically diverse organisms. First, we expressed, purified and assayed GCKs from two invertebrate species, the lancelet *Branchiostoma belcheri* and the sea squirt *Ciona intestinalis*. In the absence of added reductant, the lancelet and sea squirt enzymes displayed half-lives for inactivation of 4.6 ± 0.48 h and 7.4 ± 0.97 h, respectively (Table 1). The inclusion of 1 mM glutathione extended these values to 24.5 ± 0.78 h and 18.8 ± 0.72 h, respectively. We also expressed, purified and assayed GCKs from two vertebrates, the zebrafish *Danio rerio* and the African clawed frog *Xenopus laevis*. In the absence of glutathione, the zebrafish and frog enzymes displayed half-lives for inactivation of 1.1 ± 0.04 h and 2.6 ± 0.48 h, respectively. Like the other enzymes, 1 mM glutathione protected against inactivation and extended the half-lives to 3.8 ± 0.53 h and 7.7 ± 1.7 h, respectively. Statistical analyses of our data by Student’s t-tests demonstrate that the two invertebrate enzymes were significantly less sensitive to oxidative inactivation compared to the vertebrate GCKs (p-value < 0.05). Glutathione significantly protects all GCKs investigated against inactivation (p-value < 0.05), with comparable fold changes, ranging from 2.5 to 5.3-fold (Table 1).

### Redox sensitivity emerged during early vertebrate evolution

Our lab recently performed a comprehensive phylogenetic analysis of the glucokinase protein family using the sequences of over 190 extant enzymes.^15^ This analysis provided access to maximum likelihood reconstructions of ancestral enzymes along the vertebrate evolutionary trajectory. Characterization of the extant GCKs described above suggests that a functional transition in redox sensitivity occurred during the divergence of the invertebrate and vertebrate enzymes.^15^ To investigate this possibility further, we resurrected and functionally characterized the redox sensitivity of five ancestral enzymes representing the ancestor of chordates (cGCK), vertebrates (vGCK), gnathostomes (gGCK), tetrapods (tGCK) and mammals (mGCK). Half-lives for inactivation were determined for each enzyme using the same conditions and experimental protocol used to characterize the extant GCKs described above. As shown in Table 1, cGCK was much more resistant to oxidative inactivation compared to any extant protein we investigated, with a half-life of inactivation of ∼ 2.5 days at room temperature in the absence of any added reductant. Interestingly, the slow loss of cGCK activity observed in these studies was insensitive to the presence of glutathione, suggesting that inactivation is not associated with a redox-dependent process. By contrast, the other ancestors displayed half-lives for inactivation that were of comparable magnitude to the extant GCKs we had sampled. In addition, the vGCK, gGCK, tGCK, and mGCK enzymes displayed statistically significant protection by glutathione (p-value < 0.05), with comparable fold changes as those observed with extant enzymes.

### Cysteine content does not correlate with redox sensitivity in GCKs

Evolutionary studies focusing on the origins of redox sensitivity in other proteins suggest that the emergence of redox sensitivity coincides with an increase in overall cysteine content.^16,17^ Indeed, the abundance of cysteines is known to increase with organismal complexity.^18,19^ To investigate the extent to which differences in cysteine content correlate with our observed differences in redox sensitivity, we examined the total number of cysteine residues in extant and ancestral enzymes. Human GCK, the benchmark for the present study, contains 12 total cysteine residues. Only the zebrafish GCK contains more cysteine residues, with a total of 15. Interestingly, this protein displays a slightly increased susceptibility to oxidative inactivation compared to human GCK, both in the presence and absence of GSH (Table 1, p-value < 0.05). Frog GCK and sea squirt GCK both contain 11 cysteine residues. The frog enzyme displays a slight, but statistically significant reduction in sensitivity to oxidative inactivation in the absence of GSH (Table 1, p-value < 0.05), but not in the presence of GSH. Sea squirt GCK shows a larger increase in resistance to oxidative inactivation, despite having the same overall number of cysteines. Finally, the lancelet GCK contains the fewest cysteine residues, 10, and displays similar redox sensitivity as the sea squirt enzyme. Overall, our data do not suggest the existence of a correlation between susceptibility to oxidative inactivation and cysteine content for the extant GCKs that we sampled.

Four of the five ancestral enzymes sampled here, cGCK, vGCK, gGCK and mGCK have one additional cysteine not found in human GCK (Figure 2, blue bars). The location of this additional cysteine residue is variable and not conserved among any of these four proteins. The tetrapod ancestor has the same number and positional distribution of cysteines as human GCK, whereas the oldest ancestor, cGCK, contains one less cysteine residue. Akin to our observations with the extant enzymes, there appears to be no direct connection between total cysteine content and redox sensitivity in these ancestral GCKs.

**Figure 2.**
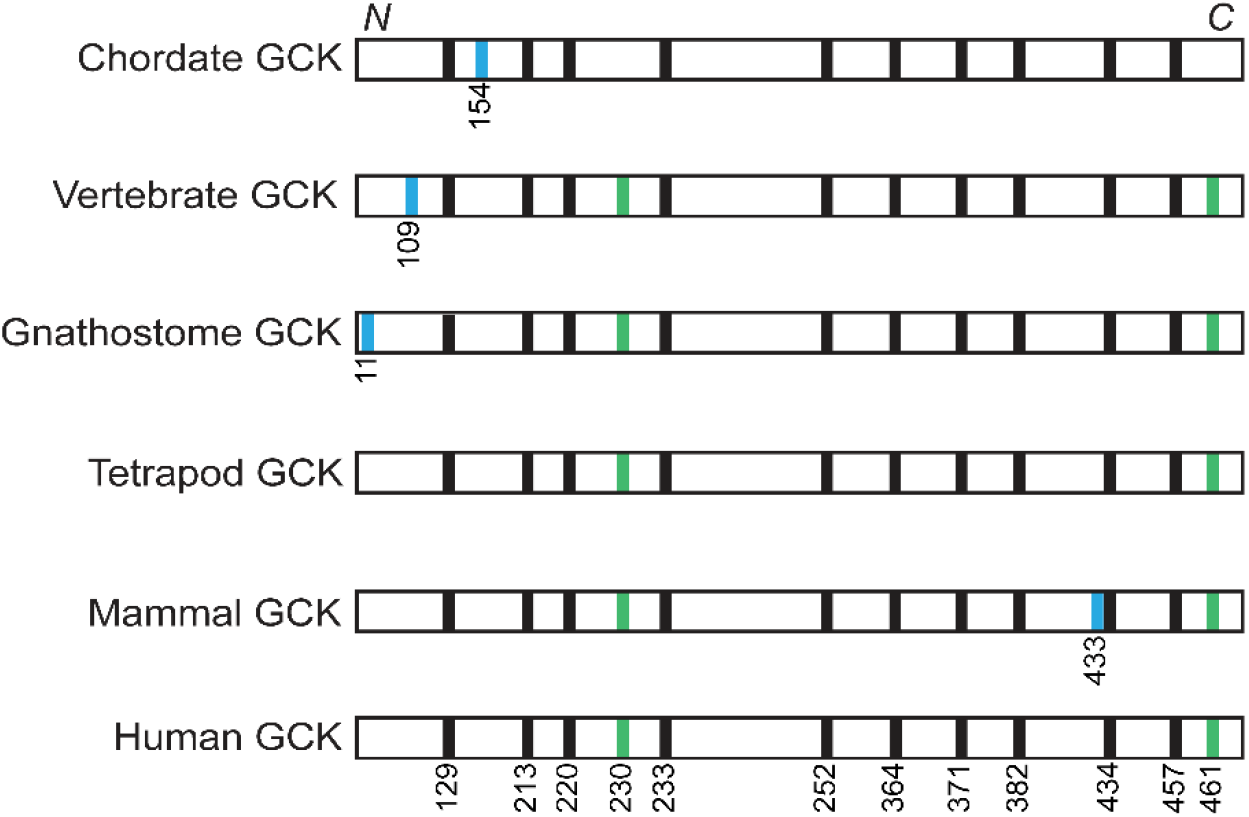
Cysteine content of ancestral GCKs and extant human GCK. Residue numbering is based on human GCK. Blue bars represent cysteine residues that are unique to individual enzymes. Green bars represent cysteine residues that are present in all redox sensitive ancestors but absent in the insensitive cGCK ancestor.

### The origins of redox sensitivity in vertebrate GCKs

Across the entire vertebrate phylogeny, which is estimated to span ∼450 million years, 10 of 12 cysteine residues found in human GCK are completely conserved in all ancestors (Figure 2, black bars). cGCK is the outlier in terms of oxidative stability. Notably, cGCK lacks two cysteine residues at positions 230 and 461, both of which are present in all other redox responsive ancestors (Figure 2, green bars). The two invertebrate GCKs, which were significantly less sensitive to oxidative inactivation compared to the vertebrate enzymes, also lack cysteine residues at these two positions. These observations raise the interesting possibility that the differences in redox sensitivity may originate from differences in cysteine content at one and/or both polypeptide positions. To investigate the extent to which these residues contribute to the redox sensitivity of vGCK, each cysteine was converted to the corresponding cGCK residue to produce the C230S and ΔC461 vGCK variants. The ΔC461 variant was created by deleting the final ten residues of the vertebrate ancestor, since the C-terminus of vGCK is ten residues longer than in cGCK (Figure S1). vGCK also contains a cysteine residue at position 109, which is not found in cGCK or any other ancestral enzymes. To investigate whether this additional cysteine residue contributes to the increase in redox sensitivity that we observed between cGCK and vGCK, we also prepared the C109R vGCK variant.

Steady-state kinetic analyses demonstrated that all three variants displayed kinetic characteristics largely unchanged from wild-type vGCK (Table 2). The biggest differences were observed in the *K*_m_ values for ATP. The C109R variant displayed a 3.9-fold reduction in this parameter and the C230S variant displayed a 2.6-fold increase. As shown in Table 3, all three variants displayed statistically significant increases in their half-lives for inactivation in the absence of glutathione compared to the wild-type vGCK ancestor. Notably, the C190R variant was least susceptible to activity loss, with a half-life of 37.1 ± 3.8 hours and this loss was independent of the presence of glutathione, suggesting that it was unrelated to oxidation. In contrast, the C230S and ΔC461 variants were partially protected against inactivation by glutathione, but only the ΔC461 variant displayed a statistically significant difference. Together these data suggest that all three positions contribute to the sensitivity of the vGCK to oxidative inactivation, and the presence of cysteine at position 109 plays the largest role.

**Table 2.** Steady-state kinetic parameters for vGCK, cGCK and variants thereof.

| | $k_{cat}$ ($s^{-1}$) | Glucose $K_{0.5}/K_m$ (mM) | Hill | ATP $K_m$ (mM) |
| --- | --- | --- | --- | --- |
| vGCK* | $54.0 \pm 6.9$ | $2.9 \pm 0.2$ | $1.6 \pm 0.1$ | $0.70 \pm 0.17$ |
| C109R vGCK | $36.0 \pm 1.1$ | $2.8 \pm 0.1$ | $1.8 \pm 0.2$ | $0.18 \pm 0.01$ |
| C230S vGCK | $32.0 \pm 1.5$ | $4.5 \pm 0.6$ | $1.5 \pm 0.2$ | $1.80 \pm 0.19$ |
| ΔC461 vGCK | $59.0 \pm 0.6$ | $3.4 \pm 0.3$ | $1.7 \pm 0.1$ | $0.44 \pm 0.01$ |
| cGCK* | $46.6 \pm 7.9$ | $0.077 \pm 0.007$ | N/A | $0.41 \pm 0.02$ |
| R109C cGCK | $38.1 \pm 2.3$ | $0.077 \pm 0.006$ | N/A | $0.71 \pm 0.07$ |
| S230C cGCK | $44.1 \pm 1.9$ | $0.077 \pm 0.004$ | N/A | $0.16 \pm 0.01$ |
| *ins*C461 vGCK | $44.5 \pm 6.3$ | $0.063 \pm 0.006$ | N/A | $0.30 \pm 0.07$ |
| R109C-S230C-*ins*C461 | $41.0 \pm 4.9$ | $0.076 \pm 0.010$ | N/A | $0.38 \pm 0.12$ |

\*from reference 15. Values are the average ± standard deviation.

**Table 3.**
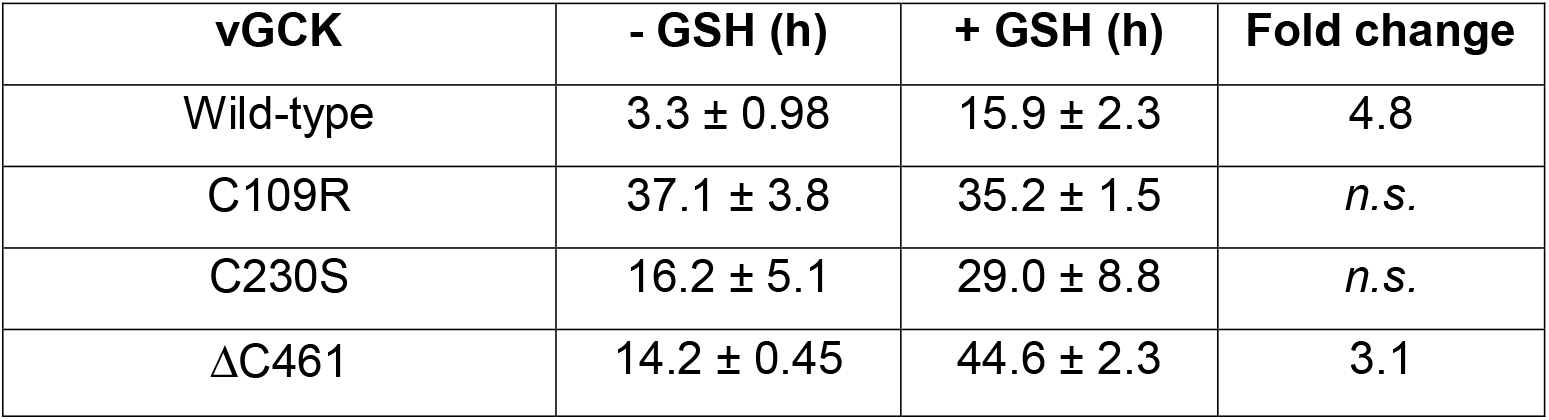
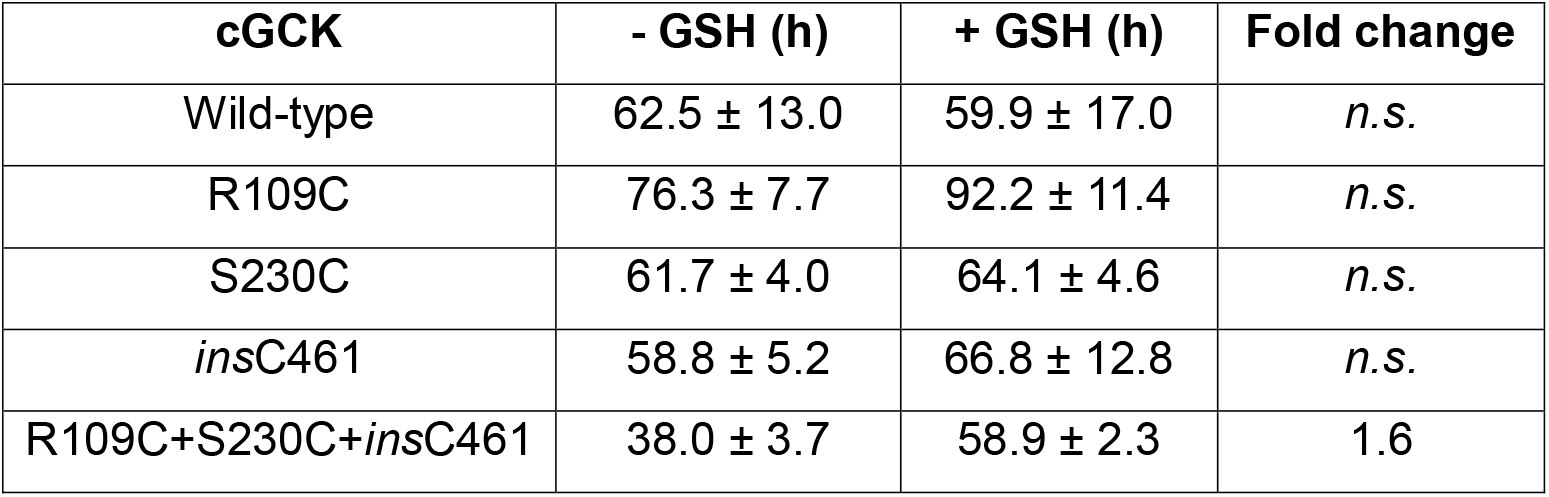
Half-lives for oxidative inactivation of vGCK and cGCK variants in the absence and presence of 1 mM glutathione (GSH). *n*.*s*. indicates no significant difference.

Next, we investigated whether insertion of cysteines at positions 109, 230 or 461 could install redox sensitivity into cGCK. To insert a cysteine residue at position 461, we appended the final ten residues of vGCK onto the C-terminus of cGCK (Figure S1). Steady-state kinetic assays indicated that the R109C, S230C and ins461 variants were functionally similar to the parental cGCK (Table 2). As shown in Table 3, all three variants were insensitive to oxidative inactivation, as evidenced by their long inactivation half-lives that were independent of glutathione. These data demonstrate that positional insertion of individual cysteine residues into cGCK did not alter the intrinsic redox insensitivity of the parental ancestor. Finally, we also prepared a cGCK triple variant that contains all three cysteine substitutions, R109C+S230C+*ins*C461. This variant also displayed kinetic characteristics indistinguishable from wild-type cGCK (Table 2). The combined insertion of all three cysteines conferred mild redox sensitivity, as reflected by a decrease in the half-live for inactivation to 38.0 ± 3.7 h, which is increased to 58.9 ± 2.3 h by the presence of 1 mM glutathione.

Despite having the exact same number and positional distribution of cysteine residues as vGCK, the R109C+S230C+*ins*C461 cGCK is characterized by an 11.5-fold larger half-life for inactivation in the absence of reductant. This discovery demonstrates that additional characteristics must contribute to the emergence of physiologically relevant redox sensitivity during the functional transition between the chordate and vertebrate ancestors. Among hGCK and its activated, disease-associated variants, hill coefficient and conformational heterogeneity are tightly coupled. That tight correlation is also seen in the evolutionary trajectory of vertebrate GCKs as prior biochemical and biophysical studies have demonstrated that cGCK and vGCK display notable structural differences.^15^ Pulsed proteolysis revealed that the redox sensitive vertebrate ancestor, vGCK, is highly sensitive to cleavage within a 30-residue mobile loop that is intrinsically disorder in hGCK.^15^ In contrast, the redox insensitive chordate GCK, cGCK, is resistant to proteolysis and the mobile loop adopts a folded β-hairpin in the crystal structure of unliganded cGCK.^15^ Comparative hydrogen-deuterium exchange mass spectrometry demonstrated that several peptides within the small domain of vGCK undergo more rapid exchange than similar regions of cGCK.^15^ Finally, biomolecular nuclear magnetic resonance studies confirmed that the small domain of vGCK samples a broader range of conformations in absence of glucose compared to cGCK.^15^ Notably, residues 109, 230 and 461, which we identified as sites of positional cysteine difference between redox sensitive and less sensitive GCKs, all reside within the small domain. In addition, these three positions show significant difference in solvent accessible surface area between the super-open and open states of hGCK (Table 4). Thus, it seems likely that cysteines at these positions become more solvent accessible because of the conformational expansion that occurred between cGCK and vGCK. Together, our results support a model in which the emergence of redox sensitivity in vertebrate GCKs resulted from a combination of structural alterations to the protein scaffold and the appearance of cysteine residues at specific positions, particularly at positions 230 and/or 461.

**Table 4.** Changes in solvent accessible surface area (SASA) of individual cysteine residues in hGCK between the open (PDB 4DCH) and super-open (PDB 1V4T) conformations computed using the solvent accessible surface area tool in PyMOL.

| Residue | SASA<sub>open</sub> (Å<sup>2</sup>) | SASA<sub>super-open</sub> (Å<sup>2</sup>) | Change (Å<sup>2</sup>) |
| --- | --- | --- | --- |
| Cys129 | 5.7 | 8.6 | +2.9 |
| Cys213 | 0.6 | 2.5 | +1.9 |
| Cys220 | 3.0 | 2.3 | -0.7 |
| Cys230 | 0.0 | 5.0 | +5.0 |
| Cys233 | 0.7 | 1.4 | +0.7 |
| Cys252 | 0.0 | 0.1 | +0.1 |
| Cys364 | 4.2 | 5.3 | +0.9 |
| Cys371 | 0.0 | 3.7 | +3.7 |
| Cys382 | 0.0 | 0.0 | — |
| Cys434 | 21 | 24 | +3.0 |
| Cys457 | 0.0 | 75 | +75 |
| Cys461 | 22 | 150 | +128 |

## Conclusions

Similar to the emergence of regulation of GCKs by allostery and by a protein-protein interaction with GKRP, the appearance of redox regulation in GCK was facilitated by conformational expansion that included access to new states. Increased protein dynamics and/or increased conformational heterogeneity have been postulated to contribute to the emergence of redox sensitivity in the adenosine 5’-phosphosulfate kinase and Aurora kinase protein families.^20,21^ Our data support a similar view and suggest that altering conformational landscapes may be a simple and common evolutionary mechanism for increasing regulatory complexity that correlates with the evolution of organismal complexity.

## ASSOCIATED CONTENT

### Supporting Information

The following Supporting Information is available free of charge online:

Protein and DNA primer sequences. Inactivation profile of hGCK. Representative Michaelis-Menten/Hill plots for all enzymes.

## ACKNOWLEDGMENTS

Research reported in this publication was supported by the National Institute of General Medical Sciences of the National Institutes of Health under Award Numbers R01GM133843 and R01GM157172 (B.G.M.) The content is solely the responsibility of the authors and does not necessarily represent the official views of the National Institutes of Health. It was also supported by grant 047082 from the FSU Council on Research and Creativity (B.G.M.), and the Pfeiffer Professorship for Cancer Research (B.G.M.).

## Notes

### Competing Interest Statement

The authors have declared no competing interest.

## REFERENCES

(1) Sternisha, S. M., and Miller, B. G. (2019) Molecular and cellular regulation of human glucokinase. Arch. Biochem. Biophys. 663, 199–213.

(2) Larion, M., and Miller, B. G. (2012) Homotropic allosteric regulation in monomeric mammalian glucokinase. Arch. Biochem. Biophys. 519, 103–111.

(3) Whittington, A. C., Larion, M., Bowler, J. M., Ramsey, K. M., Brüschweiler, R., and Miller, B. G. (2015) Dual allosteric activation mechanisms in monomeric human glucokinase. Proceedings of the National Academy of Sciences 112, 11553–11558.

(4) Kamata, K., Mitsuya, M., Nishimura, T., Eiki, J., and Nagata, Y. (2004) Structural Basis for Allosteric Regulation of the Monomeric Allosteric Enzyme Human Glucokinase. Structure 12, 429–438.

(5) Larion, M., Salinas, R. K., Bruschweiler-Li, L., Brüschweiler, R., and Miller, B. G. (2010) Direct Evidence of Conformational Heterogeneity in Human Pancreatic Glucokinase from High-Resolution Nuclear Magnetic Resonance. Biochemistry 49, 7969–7971.

(6) Beck, T., and Miller, B. G. (2013) Structural Basis for Regulation of Human Glucokinase by Glucokinase Regulatory Protein. Biochemistry 52, 6232–6239.

(7) Tippett, P. S., and Neet, K. E. (1983) Interconversions between different sulfhydryl-related kinetic states in glucokinase. Arch. Biochem. Biophys. 222, 285–298.

(8) Rizzo, M. A., and Piston, D. W. (2003) Regulation of beta cell glucokinase by S-nitrosylation and association with nitric oxide synthase. J. Cell Biol. 161, 243–8.

(9) Cullen, K. S., Matschinsky, F. M., Agius, L., and Arden, C. (2011) Susceptibility of Glucokinase-MODY Mutants to Inactivation by Oxidative Stress in Pancreatic β-Cells. Diabetes 60, 3175–3185.

(10) Tiedge, M., Richter, T., and Lenzen, S. (2000) Importance of Cysteine Residues for the Stability and Catalytic Activity of Human Pancreatic Beta Cell Glucokinase. Arch. Biochem. Biophys. 375, 251–260.

(11) Tiedge, M., Krug, U., and Lenzen, S. (1997) Modulation of human glucokinase intrinsic activity by SH reagents mirrors post-translational regulation of enzyme activity. Biochimica et Biophysica Acta (BBA) - Protein Structure and Molecular Enzymology 1337, 175–190.

(12) Lenzen, S., Freytag, S., and Panten, U. (1988) Inhibition of glucokinase by alloxan through interaction with SH groups in the sugar-binding site of the enzyme. Mol. Pharmacol. 34, 395–400.

(13) Meglasson, M. D., Burch, P. T., Berner, D. K., Najafi, H., and Matschinsky, F. M. (1986) Identification of Glucokinase as an Alloxan-Sensitive Glucose Sensor of the Pancreatic β-Cell. Diabetes 35, 1163–1173.

(14) Rizzo, M. A., Magnuson, M. A., Drain, P. F., and Piston, D. W. (2002) A Functional Link between Glucokinase Binding to Insulin Granules and Conformational Alterations in Response to Glucose and Insulin. Journal of Biological Chemistry 277, 34168–34175.

(15) Kamalaldinezabadi, S. S., Santiago, J. I., Papa, J. E., Wang, Y., Frantom, P. A., Li, H., Silver, R., Whittington, A. C., and Miller, B. G. Evolution of Protein Regulation in the Vertebrate Glucose Sensor. bioRxiv doi: 10.64898/2026.05.05.723016.

(16) Hansen, J. M., Jones, D. P., and Harris, C. (2020) The Redox Theory of Development. Antioxid. Redox Signal. 32, 715–740.

(17) Woehle, C., Dagan, T., Landan, G., Vardi, A., and Rosenwasser, S. (2017) Expansion of the redox-sensitive proteome coincides with the plastid endosymbiosis. Nat. Plants 3, 17066.

(18) Miseta, A., and Csutora, P. (2000) Relationship Between the Occurrence of Cysteine in Proteins and the Complexity of Organisms. Mol. Biol. Evol. 17, 1232–1239.

(19) Jordan, I. K., Kondrashov, F. A., Adzhubei, I. A., Wolf, Y. I., Koonin, E. V., Kondrashov, A. S., and Sunyaev, S. (2005) A universal trend of amino acid gain and loss in protein evolution. Nature 433, 633–638.

(20) Herrmann, J., Nathin, D., Lee, S. G., Sun, T., and Jez, J. M. (2015) Recapitulating the Structural Evolution of Redox Regulation in Adenosine 5′-Phosphosulfate Kinase from Cyanobacteria to Plants. Journal of Biological Chemistry 290, 24705–24714.

(21) Byrne, D. P., Shrestha, S., Galler, M., Cao, M., Daly, L. A., Campbell, A. E., Eyers, C. E., Veal, E. A., Kannan, N., and Eyers, P. A. (2020) Aurora A regulation by reversible cysteine oxidation reveals evolutionarily conserved redox control of Ser/Thr protein kinase activity. Sci. Signal. 13.

