## Supporting Information for "Conformational Expansion Underlies the Evolutionary Emergence of Redox Sensitivity in Vertebrate Glucokinases"

**Table of Contents**

**Figure S1**. Ancestral GCK protein sequences used in this study, aligned to hGCK

**Figure S2.** Extant GCK protein sequences used in this study, aligned to hGCK.

**Figure S3.** Oxidative inactivation decay profiles for hGCK in the presence of varying [GSH].

**Figure S4-S19**. Steady-state kinetic profiles for cGCK and vGCK variants fitted to either the Michealis-Menten or Hill equations.

**Table S1.** Primers used to construct cGCK and vGCK cysteine variants.

```
                  ....|....| ....|....| ....|....| ....|....| ....|....| ....|....| ....|....|
                           10         20         30         40         50         60         70
Chordate GCK      MHHHHHHGSG SMA~~~~~~~ ~LREEKVELI LDEFHLDNEE LNEIMGRMHK EMEKGLRKET NEDATVKMLP
Vertebrate GCK    MHHHHHHGSG SMLGRRSRME GRKGEKVEQI LSEFRLDKEE LEEVMRRMQR EMERGLRLET HEEASVKMLP
Gnathostome GCK   MHHHHHHGSG SMPGHRSRMD SCKMEKVEQI LSEFRLQKEE LKEVMRRMQR EMDRGLRLET HEEASVKMLP
Tetrapod GCK      MHHHHHHGSG SMLDHRSRMD SSKKEKVEQI LSEFRLQEED LKEVMRRMQR EMDRGLRLET HEEASVKMLP
Mammal GCK        MHHHHHHGSG SMLDHRARMD SSKKEKVEQI LSEFRLQEED LKKVMRRMQK EMDRGLRLET HEEASVKMLP
Human GCK         MHHHHHHGSG SMLDDRARME AAKKEKVEQI LAEFQLQEED LKKVMRRMQK EMDRGLRLET HEEASVKMLP

                  ....|....| ....|....| ....|....| ....|....| ....|....| ....|....| ....|....|
                           80         90        100        110        120        130        140
Chordate GCK      TYVRSLPDGT ESGDFLALDL GGTNFRVLLV KIKEGEELEG ERKVEMKSQI YRIPEDVMTG TGEQLFDYIA
Vertebrate GCK    TYVRSTPDGS EVGDFLALDL GGTNFRVMLV KVGEDE~~EG EWKVETKNQM YCIPEDVMTG TAEMLFDYIA
Gnathostome GCK   TYVRSTPDGS EVGDFLALDL GGTNFRVMLV KVGEDE~~EG EWKVETKHQM YSIPEDAMTG TAEMLFDYIA
Tetrapod GCK      TYVRSTPDGS EVGDFLALDL GGTNFRVMLV KVGEDE~~EG QWKVETKHQM YSIPEDAMTG TAEMLFDYIA
Mammal GCK        TYVRSTPEGS EVGDFLSLDL GGTNFRVMLV KVGEGE~~EG QWSVKTKHQM YSIPEDAMTG TAEMLFDYIS
Human GCK         TYVRSTPEGS EVGDFLSLDL GGTNFRVMLV KVGEGE~~EG QWSVKTKHQM YSIPEDAMTG TAEMLFDYIS

                  ....|....| ....|....| ....|....| ....|....| ....|....| ....|....| ....|....|
                          150        160        170        180        190        200        210
Chordate GCK      ECMADFLEKL GMKDRKLPLG FTFSFPCKQD GLDSASLITW TKGFSATGVE GKDVVKLLRD AIKRRGDFDM
Vertebrate GCK    ECIADFLDKL NMKHKKLPLG FTFSFPVKHE DLDKGILINW TKGFTATGAE GNNVVELLRD AIKRRGDFDM
Gnathostome GCK   ECISDFLDKH NMKHKKLPLG FTFSFPVRHE DIDKGILLNW TKGFKASGAE GNNVVGLLRD AIKRRGDFEM
Tetrapod GCK      ECISDFLDKH NMKHKKLPLG FTFSFPVRHE DIDKGILLNW TKGFKASGAE GNNVVGLLRD AIKRRGDFEM
Mammal GCK        ECISDFLDKH HMKHKKLPLG FTFSFPVRHE DIDKGILLNW TKGFKASGAE GNNIVGLLRD AIKRRGDFEM
Human GCK         ECISDFLDKH QMKHKKLPLG FTFSFPVRHE DIDKGILLNW TKGFKASGAE GNNVVGLLRD AIKRRGDFEM

                  ....|....| ....|....| ....|....| ....|....| ....|....| ....|....| ....|....|
                          220        230        240        250        260        270        280
Chordate GCK      DIVAVVNDTV GTMMSCAFED HDCLIGLIVG TGSNACYMEK MENVELLEGD KGEPNQMCIN MEWGAFGDDG
Vertebrate GCK    DVVAMVNDTV ATMISCYYED HNCEIGMIVG TGCNACYMEE MRNVELVEGE EG~~~RMCIN MEWGAFGDSG
Gnathostome GCK   DVVAMVNDTV ATMISCYYED HSCEVGMIVG TGCNACYMEE MRNVELVEGE EG~~~RMCVN TEWGAFGDTG
Tetrapod GCK      DVVAMVNDTV ATMISCYYED HRCEVGMIVG TGCNACYMEE MRNVELVEGE EG~~~RMCVN TEWGAFGDTG
Mammal GCK        DVVAMVNDTV ATMISCYYED RQCEVGMIVG TGCNACYMEE MHNVELVEGD EG~~~RMCVN TEWGAFGDAG
Human GCK         DVVAMVNDTV ATMISCYYED HQCEVGMIVG TGCNACYMEE MQNVELVEGD EG~~~RMCVN TEWGAFGDSG

                  ....|....| ....|....| ....|....| ....|....| ....|....| ....|....| ....|....|
                          290        300        310        320        330        340        350
Chordate GCK      ALDDFRTEYD REVDENSLNP GQQLYEKMIS GMYMGELVRL VLLKLTKEGL LFGGKTSEEL KTPGTFQTKY
Vertebrate GCK    ELEEFRLEYD RKVDETSLNP GQQLYEKIIS GKYMGELVRL VLLKLTNEGL LFGGKASEKL KTRGSFETKY
Gnathostome GCK   ELEEFRLEYD RVVDETSLNP GQQLYEKIIS GKYMGELVRL VLLKLVNENL LFNGEASEIL KTRGSFETRF
Tetrapod GCK      ELEEFRLEYD RVVDETSLNP GQQLYEKIIG GKYMGELVRL VLLKLVNENL LFNGEASEKL KTRGAFETRF
Mammal GCK        ELDEFLLEYD RVVDETSLNP GQQLYEKIIG GKYMGEIVRL VLLKLVDENL LFNGEASEQL RTRGAFETRF
Human GCK         ELDEFLLEYD RLVDESSANP GQQLYEKLIG GKYMGELVRL VLLRLVDENL PFHGEASEQL RTRGAFETRF

                  ....|....| ....|....| ....|....| ....|....| ....|....| ....|....| ....|....|
                          360        370        380        390        400        410        420
Chordate GCK      VSQIESDVPG DMTATLNILA SLGLRHATEV DCEIVRQVCR AVSTRAAHLC AAGIAAVVNK MRRNR~~~~~
Vertebrate GCK    VSQIESDDSG DMKQTYNILT TLGLQHPTEL DCEIVRRVCQ AVSTRAAHLC AAGMAAVVNK MRENRSQETL
Gnathostome GCK   VSQIESD~SG DRKQIYNILT TLGVLHPSEL DCDIVRLVCE SVSTRAAHMC AAGLAGVINR MRENRSQETL
Tetrapod GCK      VSQIESD~SG DRKQIYNILT TLGVLHPSAT DCDIVRLVCE SVSTRAAHMC SAGLAGVINR MRESRSEETL
Mammal GCK        VSQVESD~SG DRKQIYNILS TLGLLHPSAT DCDIVRMVCE SVSTRAAQMC SAGLAGVINR MRESRSEEVM
Human GCK         VSQVESD~TG DRKQIYNILS TLG~LRPSTT DCDIVRRACE SVSTRAAHMC SAGLAGVINR MRESRSEDVM

                  ....|....| ....|....| ....|....| ....|....| ....|....| ....|....| ....
                          430        440        450        460        470        480
Chordate GCK      ~ITVGVDGSV YKYHPTFKEL MSETVDELTP GCDVKFMLSE DGSGKGAALI TAVACRLAGK ~~~~
Vertebrate GCK    KITVGVDGSV YKLHPSFKDK FHAMVRELTP RCDITFIQSE EGSGRGAALI SAVACKMACM GQ~~
Gnathostome GCK   KITVGVDGSV YKLHPSFKDK FHAMVRELTP RCDITFIQSE EGSGRGAALI SAVACKMACM LSQ~
Tetrapod GCK      KITVGVDGSV YKLHPSFKDK FHATVRQLTP GCDITFIQSE EGSGRGAALI SAVACKMACM IGQ~
Mammal GCK        RITVGVDGSV YKLHPSFKER FHATVRQLTP CCDITFIQSE EGSGRGAALV SAVACKKACM LGQ~
Human GCK         RITVGVDGSV YKLHPSFKER FHASVRRLTP SCEITFIESE EGSGRGAALV SAVACKKACM LGQ*
```

**Figure S1.** Ancestral GCK protein sequences used in this study, aligned to hGCK.

```
B. Belcheri GCK   NPDATVKMLA TYVRAVPDGT ESGDFLALDL GGSNFRVLHV HIEK-----G AKDVRLQSDS YAIPESIMTG
C. Milii GCK      HEAASVKMLP TYVCSTPDGS EVGDFLSLDL GGTNFRVMLV KVGENEE--G DWKIATIPQM YSIPVDVTTG
D. Rerio GCK      HDEASVKMLP TYVRSTPEGS EVGDFLALDL GGTNFRVMLV KVGEDEE--R GWKVETKHHM YSIPEDAMTG
X. Laevis GCK     NEEASVKMLP TYVRSTPDGS EVGDFLALDL GGTNFRVMLV KVGEDLE--G QWKVETKHKM YSIPVDAMTG
H. sapiens GCK    HEEASVKMLP TYVRSTPEGS EVGDFLSLDL GGTNFRVMLV KVGEGEE--G QWSVKTKHQM YSIPEDAMTG

                  ....|....| ....|....| ....|....| ....|....| ....|....| ....|....| ....|....|
                          150        160        170        180        190        200        210
C. Intestinalis   KGDELFDHIA LCMADFLKKL DLLDHKLPVG FTFSFPCKQD GLDHATLITW TKGFSASNVV GKDIVKMLKT
B. Belcheri GCK   SGEQLFDYIA DCMAKFLEKI GMKEKKMALG FTFSFPCKQR GLDCASLIKW TKGFSAAGVE GEDVVRLLRD
C. Milii GCK      TAQMLFDYIA QCISNFLDRH NIKHKKLPLG FTFSFPVRHE DIDKGILLNW TKGFKASGAE GNNIVGLLRD
D. Rerio GCK      TAEMLFDYIA GCISDFLDKH NLKHKKLPLG FTFSFPVRHE DLDKGILLNW TKGFKASGAE GNNVVGLLRD
X. Laevis GCK     TAEMLFDYIA ECISDYLDQQ NMKHKKLPLG FTFSFPVRHE DIDKGILLNW TKGFKASGAE GNNVVGLLRD
H. sapiens GCK    TAEMLFDYIS ECISDFLDKH QMKHKKLPLG FTFSFPVRHE DIDKGILLNW TKGFKASGAE GNNVVGLLRD

                  ....|....| ....|....| ....|....| ....|....| ....|....| ....|....| ....|....|
                          220        230        240        250        260        270        280
C. Intestinalis   AIDKRGDLDV DIIAVVNDTV GTMTSCAFDD QECMIGLIVG TGSNACYMEK MSNIERLDSN KG---GMCIN
B. Belcheri GCK   AIKRRGDFDT DVVAVVNDTV GTMMACGLAD HDCLIGLIVG TGSNACYMEK LDNVEIWEGE RGEPNQVVVN
C. Milii GCK      AIKRRGDIEM NVVAMVNDTV ATMISSYYED HSCEVGLIVG TGCNACYMEE MKNMELVDGE EG---RMCVN
D. Rerio GCK      AIKRRGDFEM DVVAMVNDTV ATMISCYYED RSCEVGMIVG TGCNACYMEE MRKVELVEGE EG---RMCVN
X. Laevis GCK     AIKRRGDFEM DVVAMVNDTV ATMISCYYED HHCEVGLIVG TGCNACYMEE MSNVELVEGE EG---RMCVN
H. sapiens GCK    AIKRRGDFEM DVVAMVNDTV ATMISCYYED HQCEVGMIVG TGCNACYMEE MQNVELVEGD EG---RMCVN

                  ....|....| ....|....| ....|....| ....|....| ....|....| ....|....| ....|....|
                          290        300        310        320        330        340        350
C. Intestinalis   MEWGAFGDDG ALSDYRNEYD VHVDENSLNS GKQLYEKMIS GMYMGEIVRL VLVKLTEDGL LFGGETSEAL
B. Belcheri GCK   MEWGAFGEDG ALDDLRTPYD REIDEHSINR GQQIYEKMIS GMYMGELVRL VLLEMTKQML VFGGRTSTDL
C. Milii GCK      TEWGGFGDTG ELEEFRLEYD RVVDEASLNP GHQLYEKIIG GKYMGEVTRL VLLKLVNQNL LFNGEASELI
D. Rerio GCK      TEWGAFGDHS ELEDFRLEYD RVIDETSLNP GHQLYEKLIG GKYMGELVRL VLLKLVNEDL LFNGEASDLL
X. Laevis GCK     TEWGAFGDTG ELEDFRLEYD RVVDEASLNP GQQLYEKMIG GKYMGELVRL VLIKMVNENL LFGGESSEKL
H. sapiens GCK    TEWGAFGDSG ELDEFLLEYD RLVDESSANP GQQLYEKLIG GKYMGELVRL VLLRLVDENL PFHGEASEQL

                  ....|....| ....|....| ....|....| ....|....| ....|....| ....|....| ....|....|
                          360        370        380        390        400        410        420
C. Intestinalis   KTPGTFQTSY VSQIESAIPS GMTAVQNILA NLGIG-AMRA DCEVVIQVCK AVSRRAAHLC AAGIAAVARK
B. Belcheri GCK   ETRGTFQTKY VSEIEEEPIG DKNPTMKILH SLGLKHATET DCETVKEVCR AVSTRAAHLV SAGISAIVNK
C. Milii GCK      KTRGSFETKF LSQIESD-GS DQKQIYNILT TLGVL-PSEA DCEIVHLVCE SVSTRAAHIC AAGLAGVINR
D. Rerio GCK      KTRGAFETRF VSQIESD-TG DRKQIYNILS CLGIL-PSEL DCDIVRLACE SVSTRAAHLC GAGLAGVINL
X. Laevis GCK     KTRGAFETQF VSQIEAD-TS DFKQTLNILR TLGVQ-ATIG DCHAVRLACE SVSTRAAIMC SAGLAGILNR
H. sapiens GCK    RTRGAFETRF VSQVESD-TG DRKQIYNILS TLGLR-PSTT DCDIVRRACE SVSTRAAHMC SAGLAGVINR

                  ....|....| ....|....| ....|....| ....|....| ....|....| ....|....| ....|....|
                          430        440        450        460        470        480        490
C. Intestinalis   IKANHPDRET LRMTVGVDGT VYKKHPTFSQ MMSEKVDELC AGAGVDVHFA LSYDGSGKGA ALITAVAQR-
B. Belcheri GCK   MGRER----- --VAVGVDGS VYKYHPHFKE LMTQTIDKLT ---HCDVKLM LSEDGSGKGA ALVTAVACR-
C. Milii GCK      LCDNKRK-ET MKMAVGIDGS VYKLHPHFKS RFHSMVRELT --PHCDIIFL QSDEGSGRGA ALISAIACK-
D. Rerio GCK      MRERRCQ-EE LKITVGVDGS VYKLHPHFKE RFHKLVWELT --PHCEITFI QSEEGSGRGA ALISAVACKM
X. Laevis GCK     MRQSRRE-EL LRITVGVDGS VYKLHPSFKD KFHATVLKLT --SGCEITFI QSEEGSGRGA ALISAVAYK-
H. sapiens GCK    MRESRSE-DV MRITVGVDGS VYKLHPSFKE RFHASVRRLT --PSCEITFI ESEEGSGRGA ALVSAVACK-

                  ....|....

C. Intestinalis   ---KIGSDE
B. Belcheri GCK   ---LAGKS.
C. Milii GCK      ---MAT...
D. Rerio GCK      AACMLTP..
X. Laevis GCK     MAVMIGH..
H. sapiens GCK    KACMLGQ*.
```

**Figure S2.** Extant GCK protein sequences used in this study, aligned to hGCK.

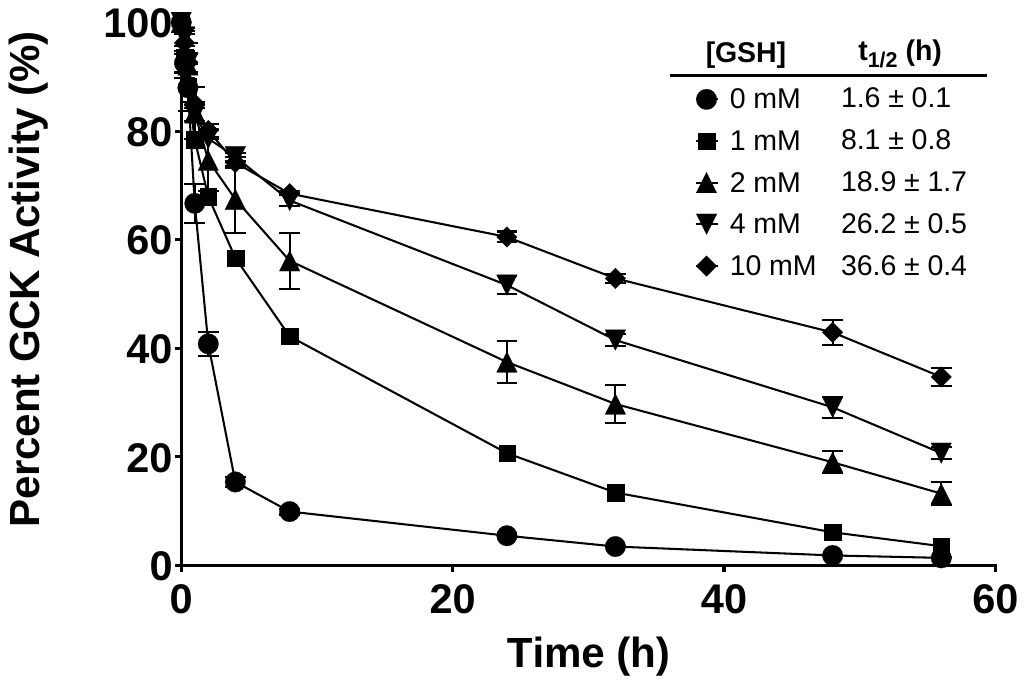

**Figure S3.** Oxidative inactivation decay profiles for hGCK in the presence of varying [GSH]. hGCK was incubated with [GSH] from 0 mM to 10 mM, and enzyme activity was monitored over 56 hours via G6PDH coupled assay. Generated time course curves were fit to a single exponential decay model, %A = %A_0_e^-kt^. The decay rate constant, *k*, was used to calculate the half-life (t_1/2_). The half-life for each [GSH] is displayed in the inset on the graph. Data displayed is from one biological replicate with technical replicates of n= 2 for [GSH] 0 and 1 mM, n=3 for [GSH] 2 to 10 mM.

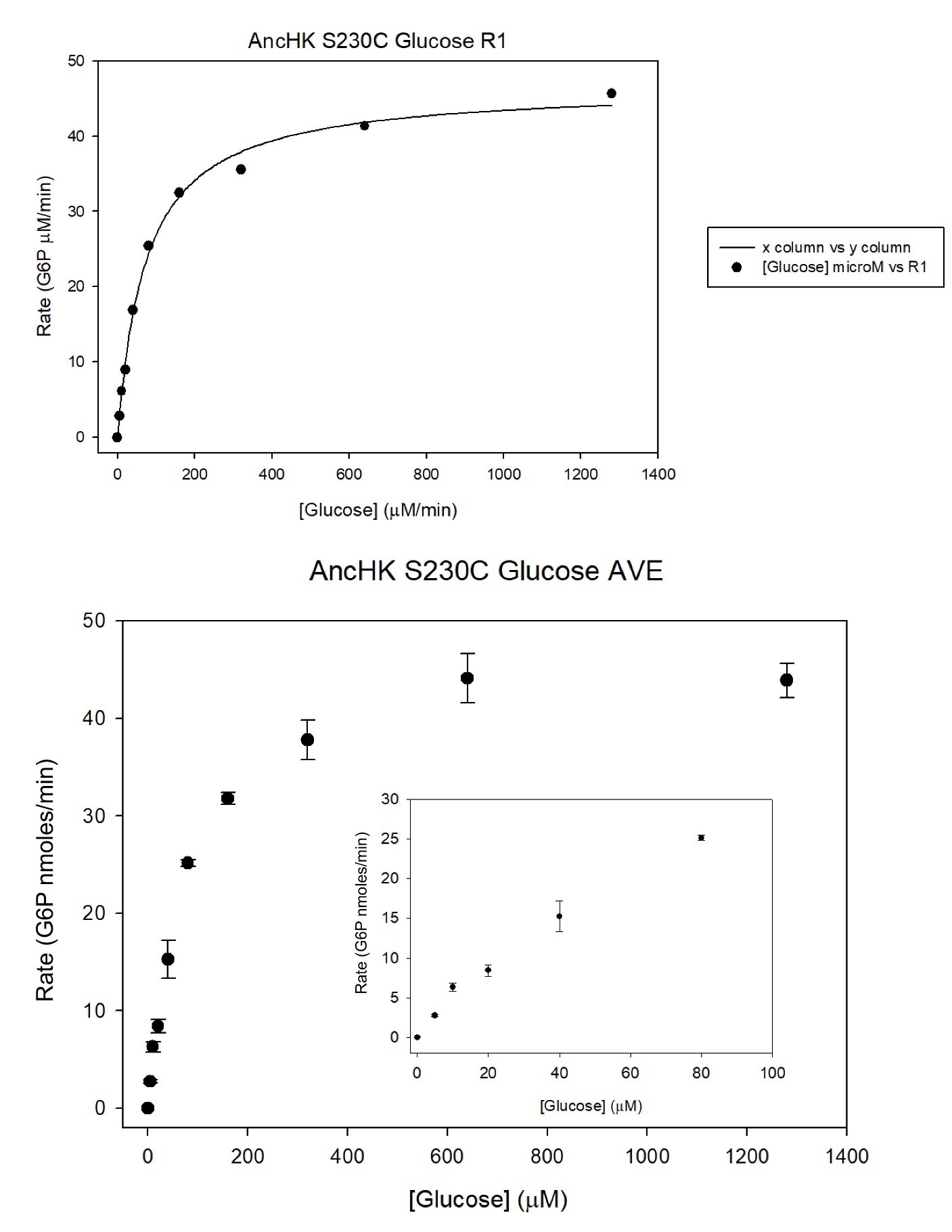

**Figure S4.** Steady-state glucose kinetic profile for S230C cGCK fitted to Michaelis-Menten equation, with n = 3 technical replicates. Inset is the linear portion of curve. *K_M_* = 0.08 ± 0.01 mM, *k_cat_* = 47.2 ± 1.4 s^-1^, (Ave ± SD).

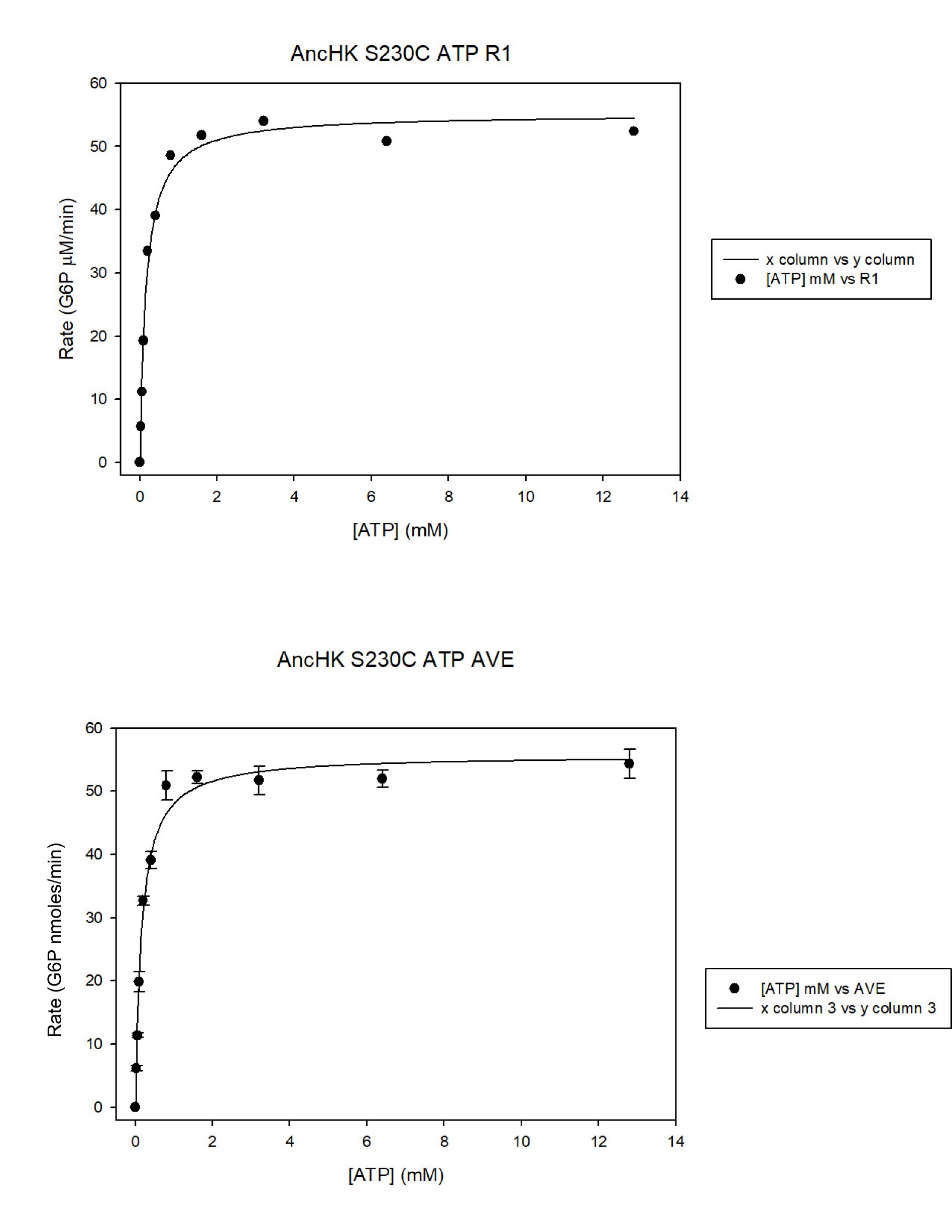

**Figure S5.** Steady-state ATP kinetic profile for S230C cGCK fitted to Michaelis-Menten equation, with n = 3 technical replicates. *K_M_* = 0.16 ± 0.01 mM, *k_cat_* = 46.4 ± 0.6 s^-1^, (Ave ± SD).

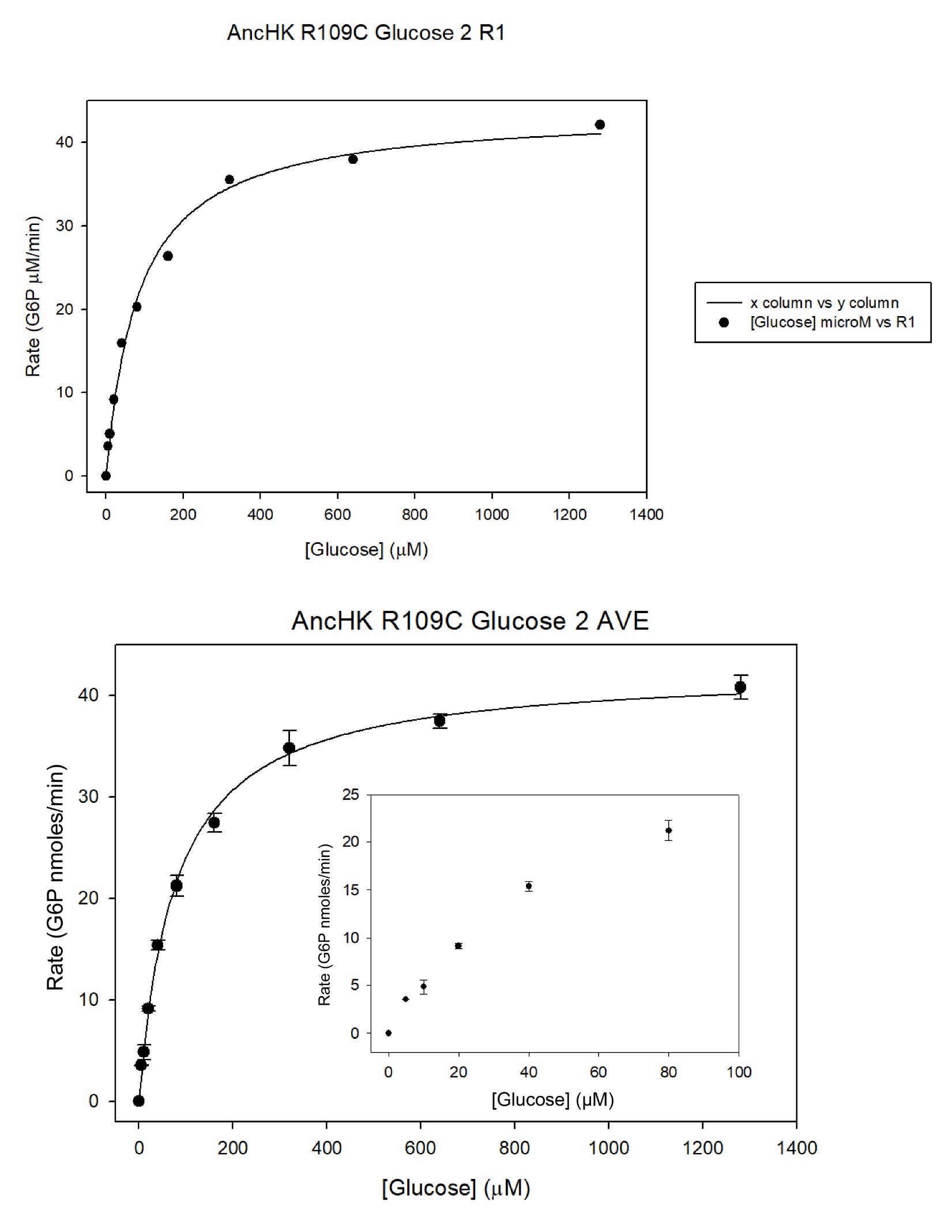

**Figure S6.** Steady-state glucose kinetic profile for R109C cGCK fitted to Michaelis-Menten equation, with n = 3 technical replicates. Inset is the linear portion of curve. *K_M_* = 0.077 ± 0.006 mM, *k_cat_* = 36.5 ± 1.1 s^-1^, (Ave ± SD).

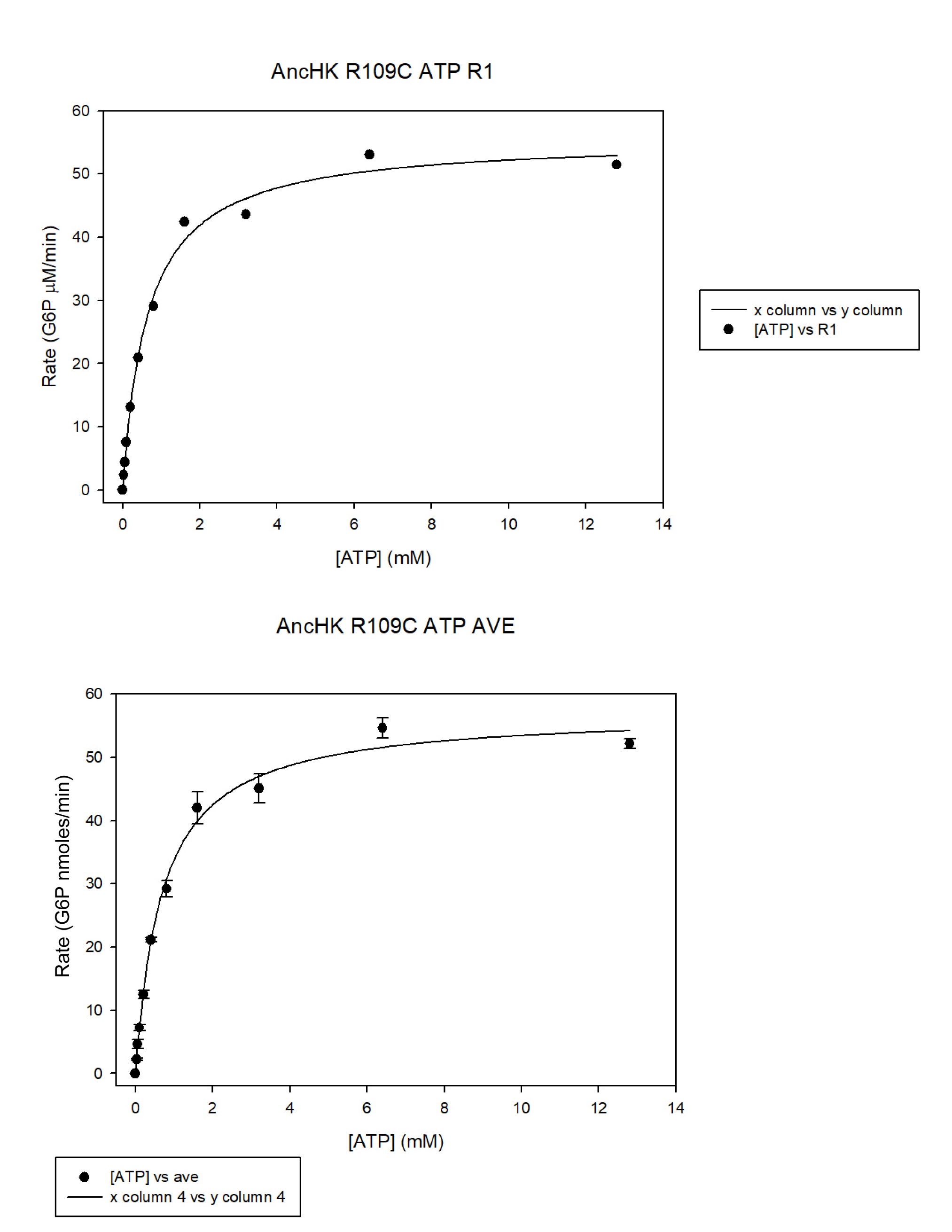

**Figure S7.** Steady-state ATP kinetic profile for R109C cGCK fitted to Michaelis-Menten equation, with n = 3 technical replicates. *K_M_* = 0.71 ± 0.07 mM, *k_cat_* = 49.1 ± 1.9 s^-1^, (Ave ± SD).

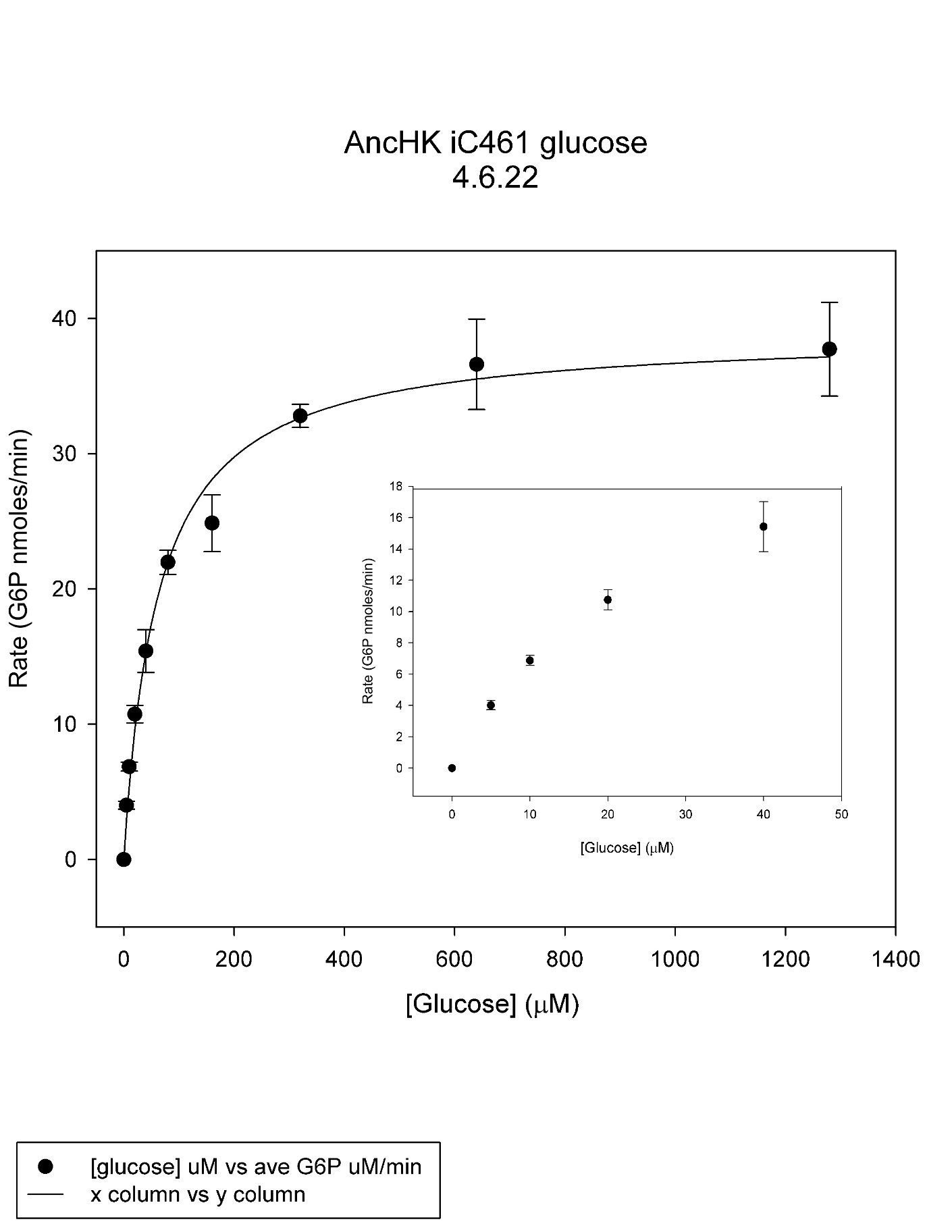

**Figure S8.** Steady-state glucose kinetic profile for InsC461 cGCK fitted to Michaelis-Menten equation, with n = 3 technical replicates. Inset is the linear portion of curve. *K_M_* = 0.059 ± 0.003 mM, *k_cat_* = 43.3 ± 3.4 s^-1^, (Ave ± SD).

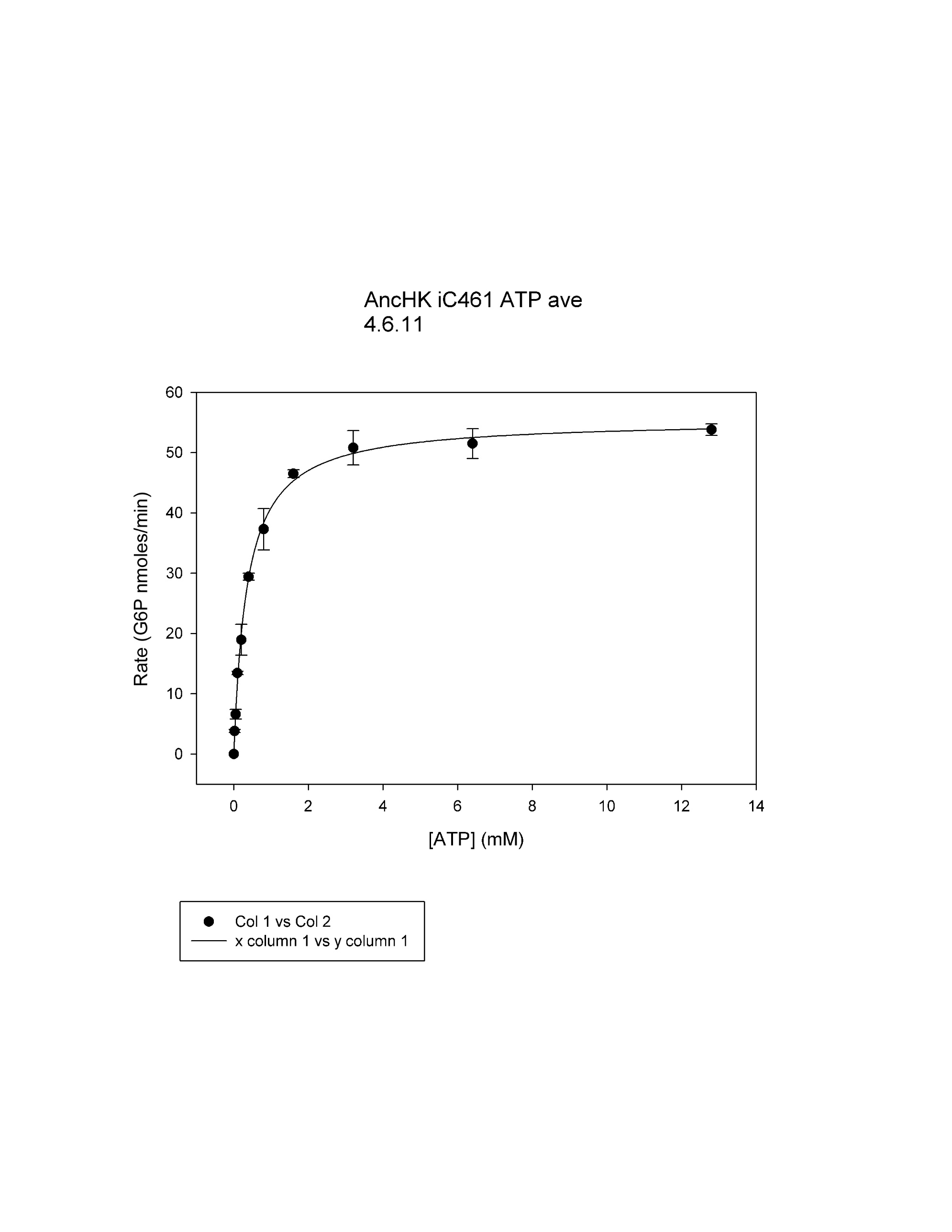

**Figure S9.** Steady-state ATP kinetic profile for InsC461 cGCK fitted to Michaelis-Menten equation, with n = 3 technical replicates. *K_M_* = 0.0.36 ± 0.02 mM, *k_cat_* = 61.4 ± 1.2 s^-1^, (Ave ± SD).

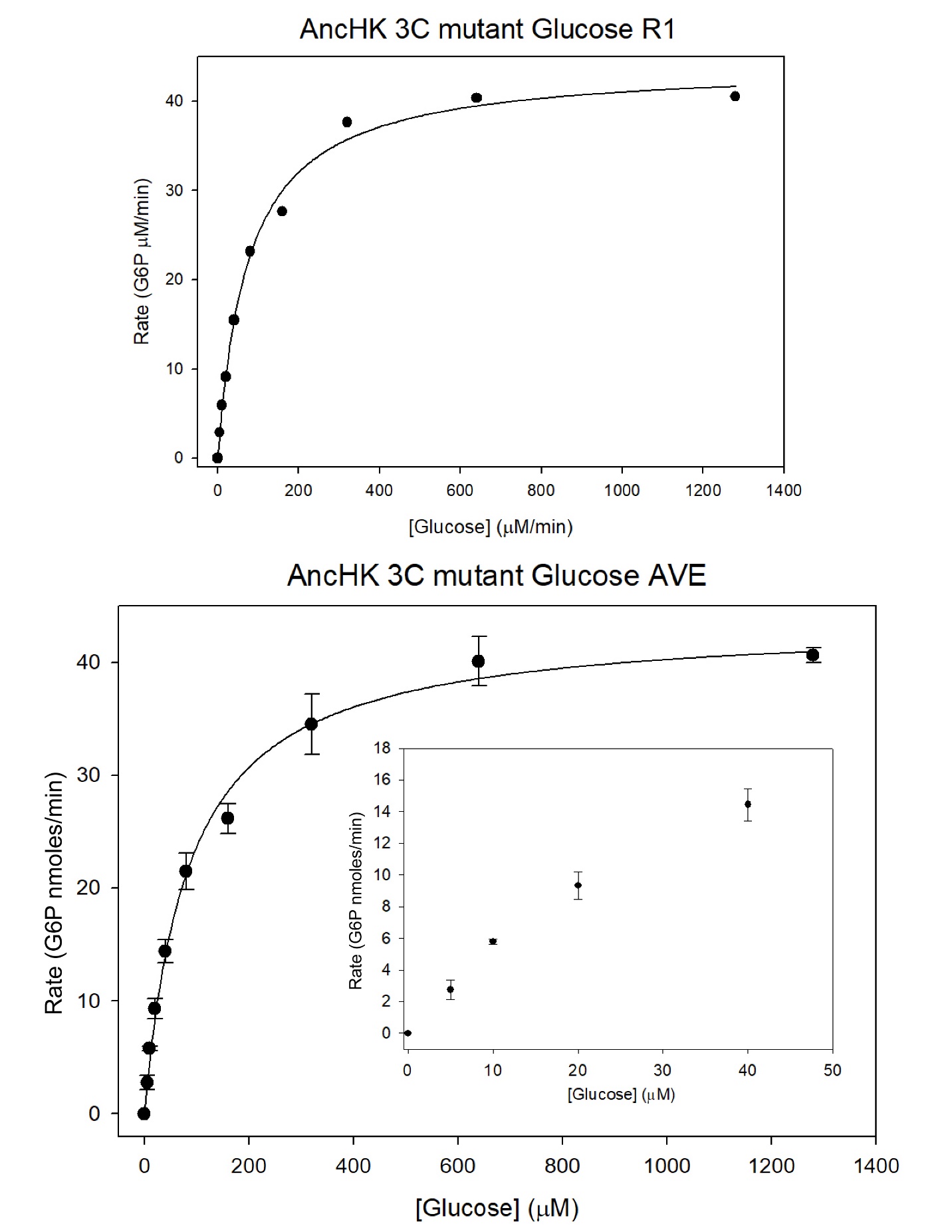

**Figure S10.** Steady-state glucose kinetic profile for R109C-S230C-InsC461 cGCK fitted to Michaelis-Menten equation, with n = 3 technical replicates. Inset is the linear portion of curve. *K_M_* = 0.053 ± 0.008 mM, *k_cat_* = 32.7 ± 1.0 s^-1^, (Ave ± SD).

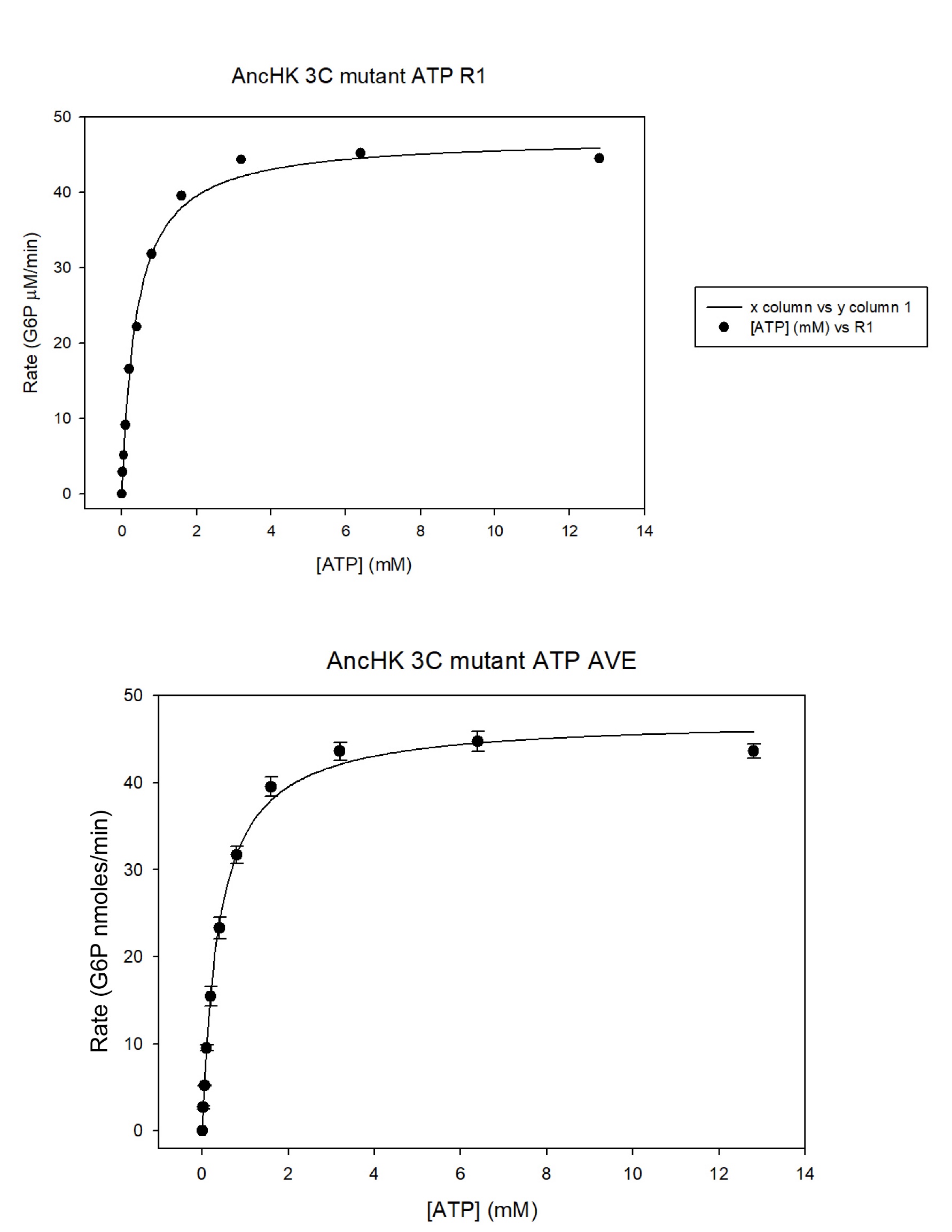

**Figure S11.** Steady-state ATP kinetic profile for R109C-S230C-InsC461 cGCK fitted to Michaelis-Menten equation, with n = 3 technical replicates. *K_M_* = 0.15 ± 0.002 mM, *k_cat_* = 34.8 ± 0.62 s^-1^, (Ave ± SD).

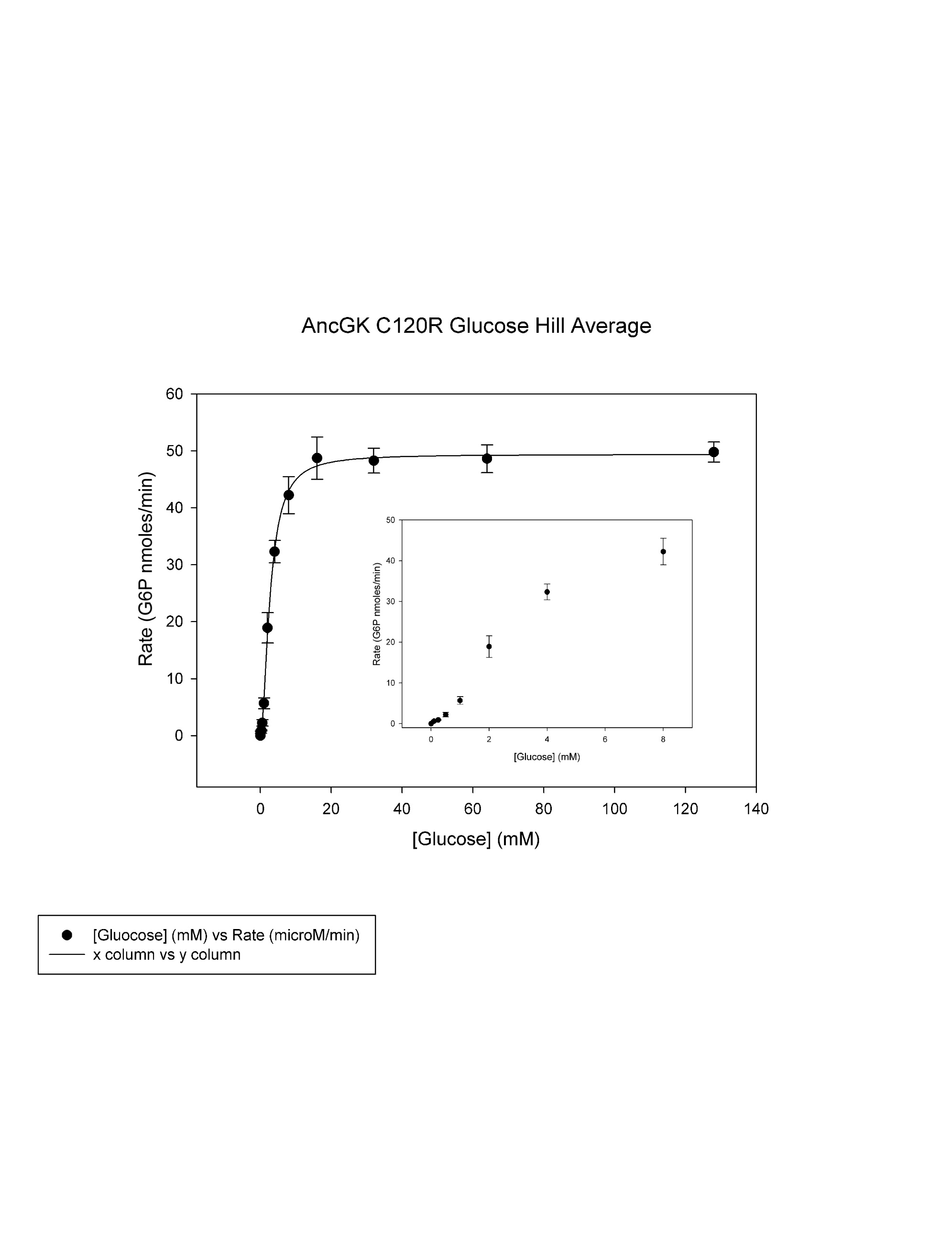
**Figure S12.** Steady-state glucose kinetic profile for C109R vGCK fitted to Hill equation, with n = 3 technical replicates. Inset is the sigmoidal portion of curve. *K_0.5_* = 2.8 ± 0.1 mM, *k_cat_* = 36.8 ± 1.1 s^-1^, hill =1.8 ± 0.3, (Ave ± SD).

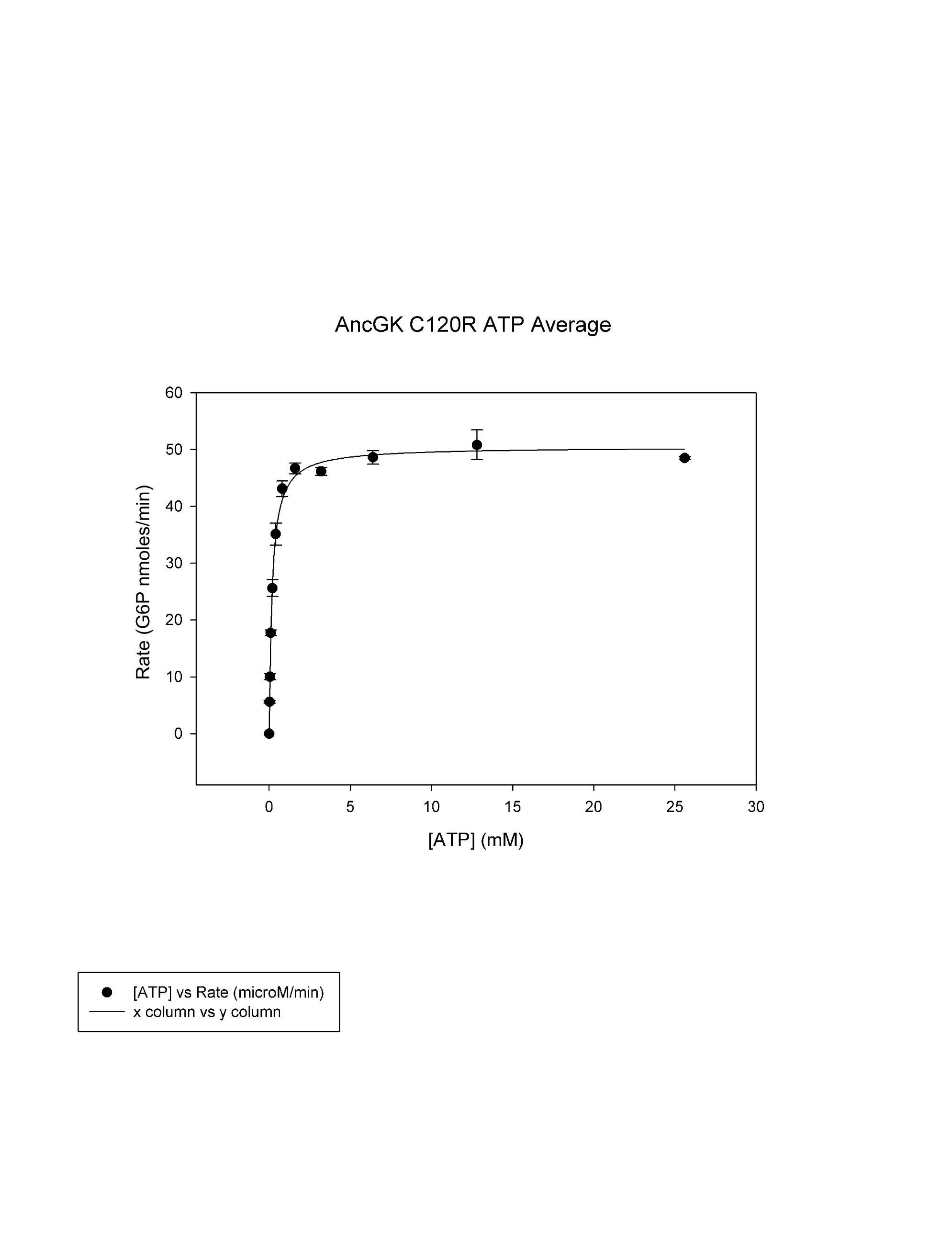

**Figure S13.** Steady-state ATP kinetic profile for C109R vGCK fitted to Michaelis-Menten equation, with n = 3 technical replicates. *K_M_* = 0.18 ± 0.01 mM, *k_cat_* = 38.2 ± 0.6 s^-1^, (Ave ± SD).

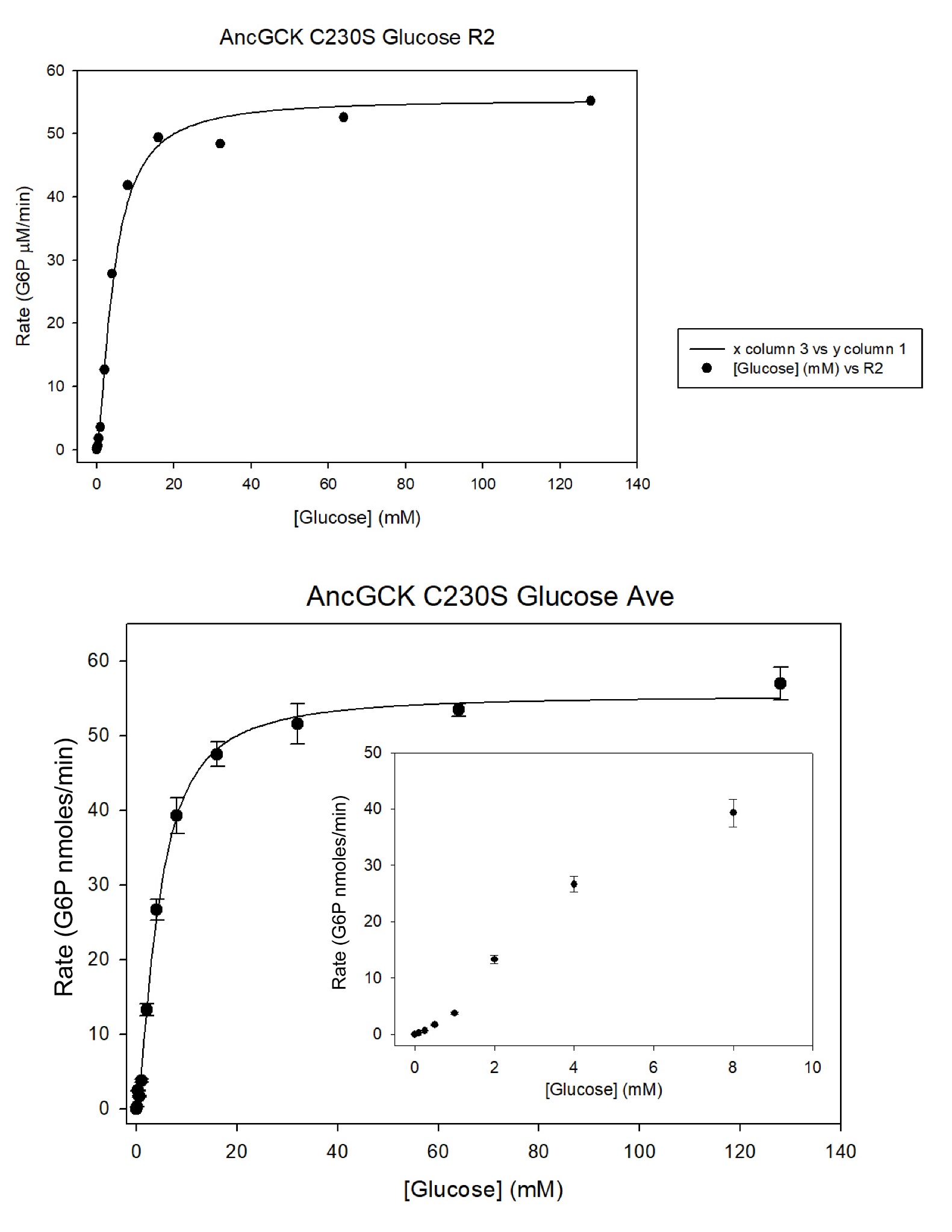

**Figure S14.** Steady-state glucose kinetic profile for C230S vGCK fitted to Hill equation, with n = 3 technical replicates. Inset is the sigmoidal portion of curve. *K_0.5_* = 4.5 ± 0.6 mM, *k_cat_* = 33 ± 1.5 (s^-1^), hill =1.5 ± 0.3, (Ave ± SD).

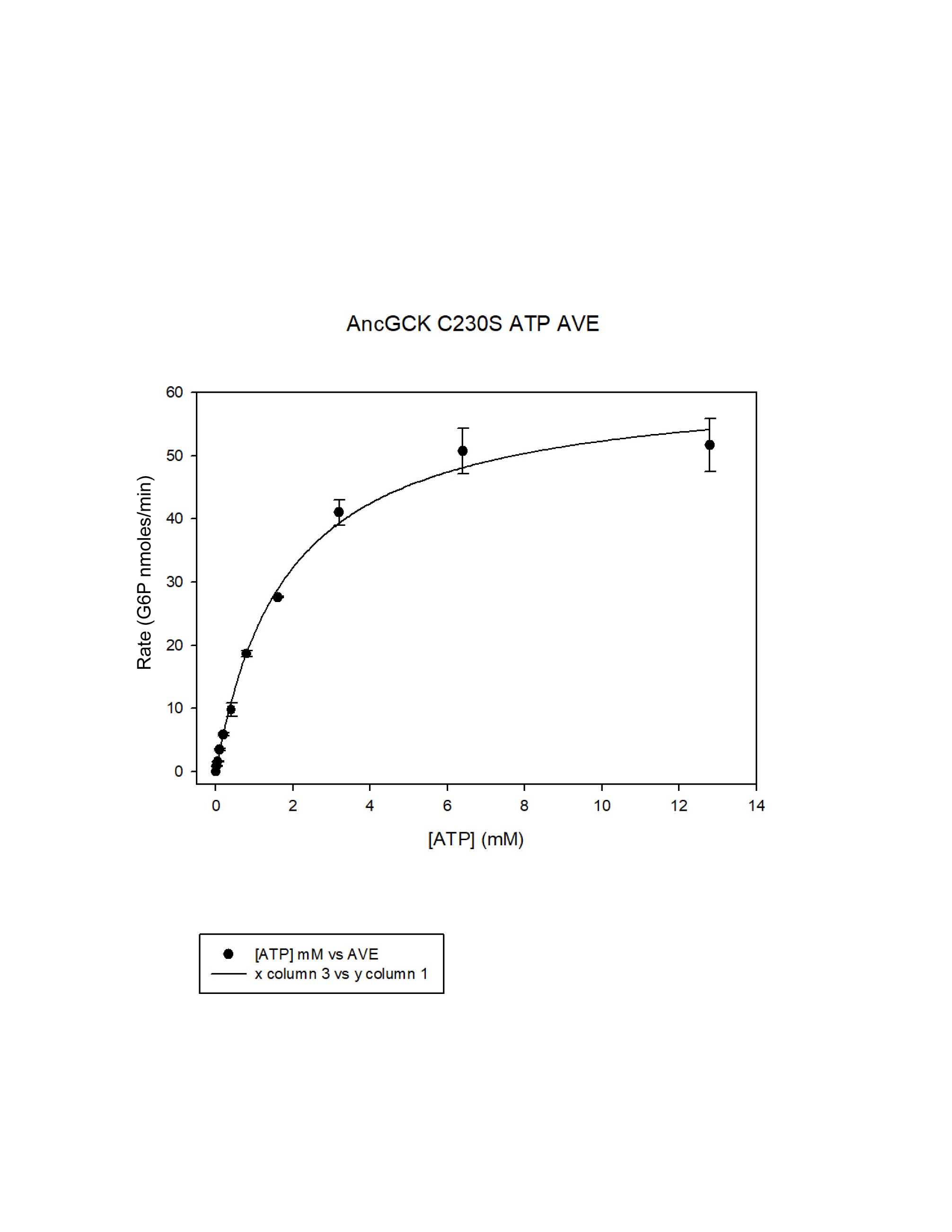

**Figure S15.** Steady-state ATP kinetic profile for C230S vGCK fitted to Michaelis-Menten equation, with n = 3 technical replicates. *K_M_* = 1.8 ± 0.2 mM, *k_cat_* = 37.0 ± 2.2 s^-1^, (Ave ± SD).

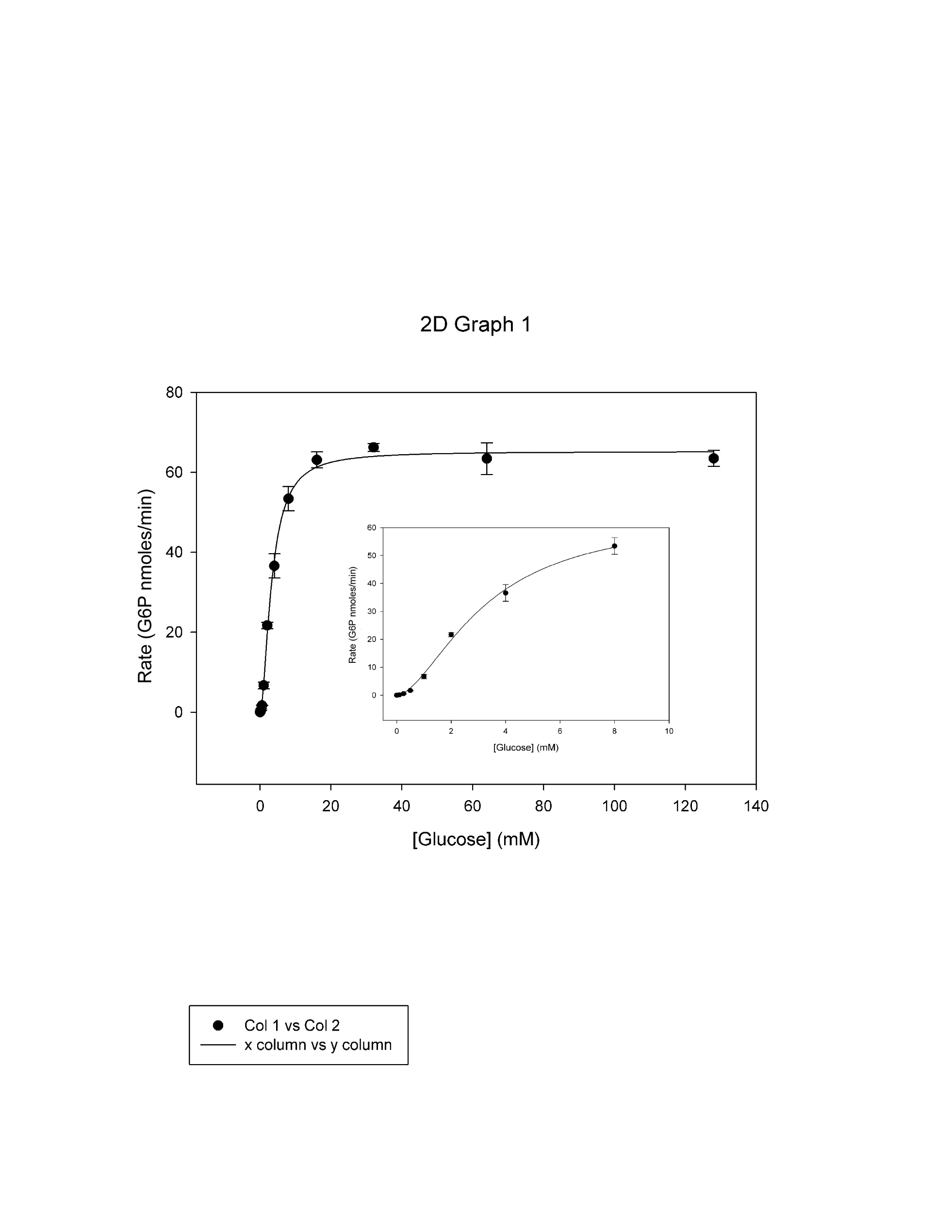

**Figure S16.** Steady-state glucose kinetic profile for ΔC461 vGCK fitted to Hill equation, with n = 3 technical replicates. Inset is the sigmoidal portion of curve. *K_0.5_* = 3.4 ± 0.3 mM, *k_cat_* = 41.9 ± 2.2 (s^-1^), hill =1.7 ± 0.2, (Ave ± SD).

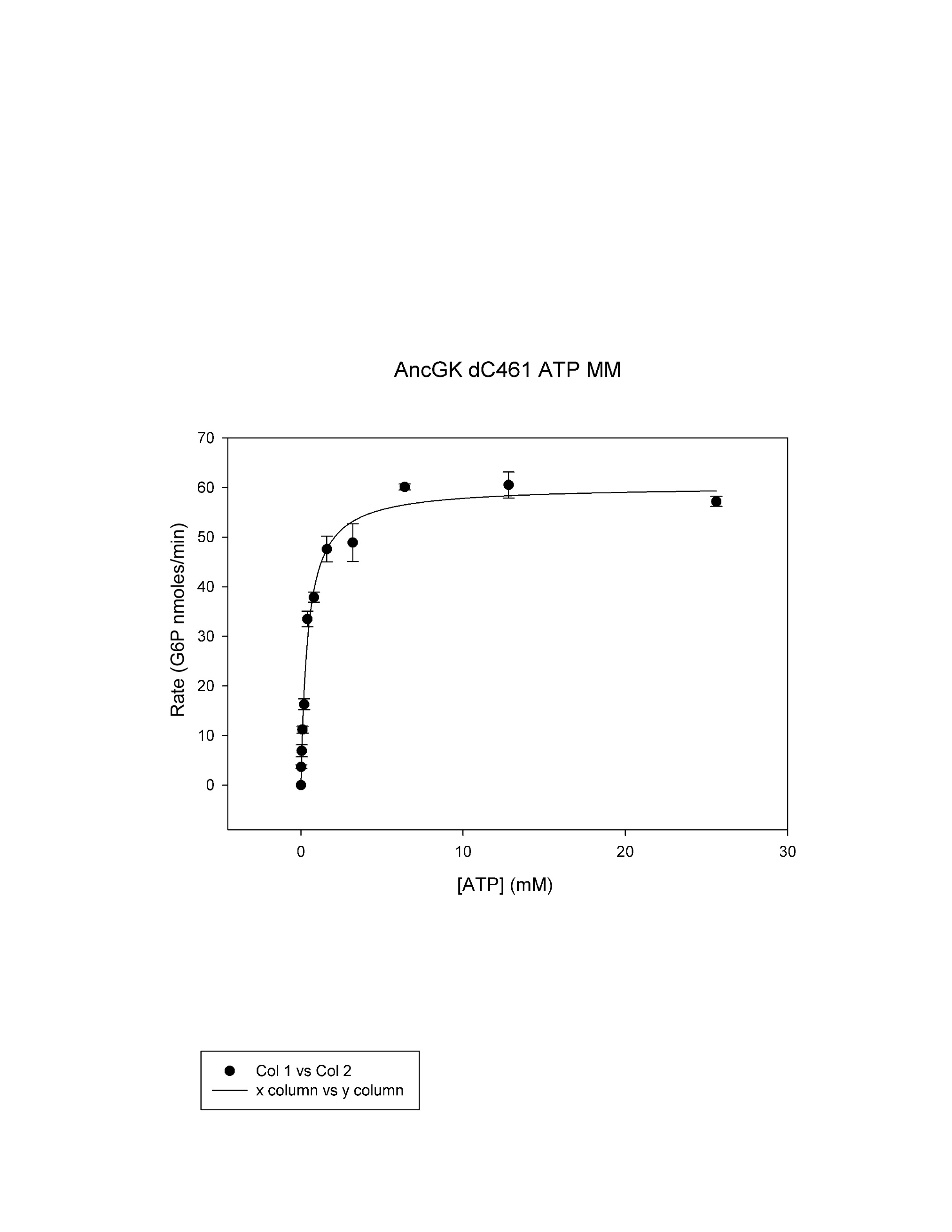

**Figure S17.** Steady-state ATP kinetic profile for ΔC461 vGCK fitted to Michaelis-Menten equation, with n = 3 technical replicates. *K_M_* = 0.44 ± 0.03 mM, *k_cat_* = 39.4 ± 0.3 s^-1^, (Ave ± SD).

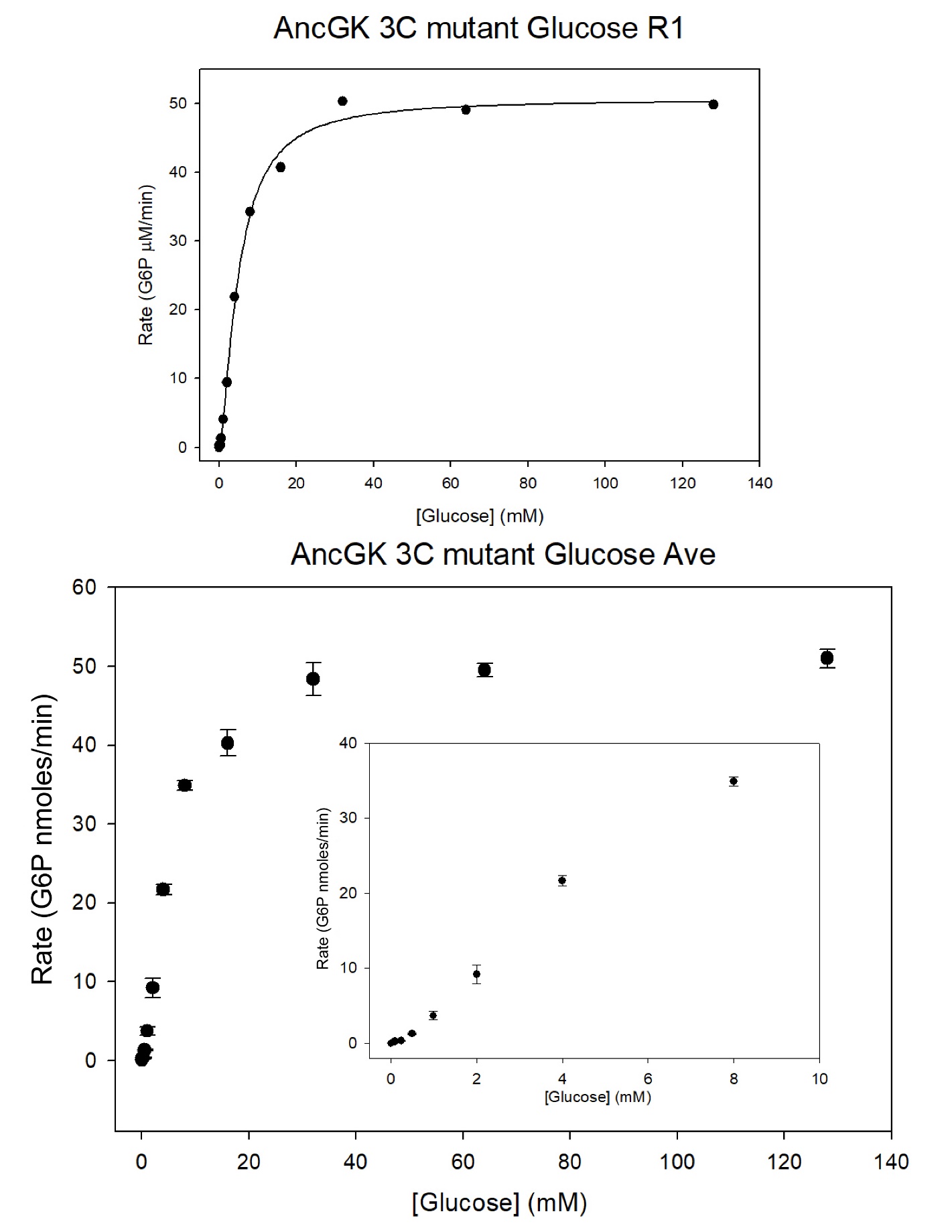

**Figure S18.** Steady-state glucose kinetic profile for C109R+C230S+ΔC461 vGCK fitted to Hill equation, with n = 3 technical replicates. Inset is the sigmoidal portion of curve. *K_0.5_* = 5.1 ± 0.3 mM, *k_cat_* = 35.0 ± 0.2 (s^-1^), hill =1.5 ± 0.05, (Ave ± SD).

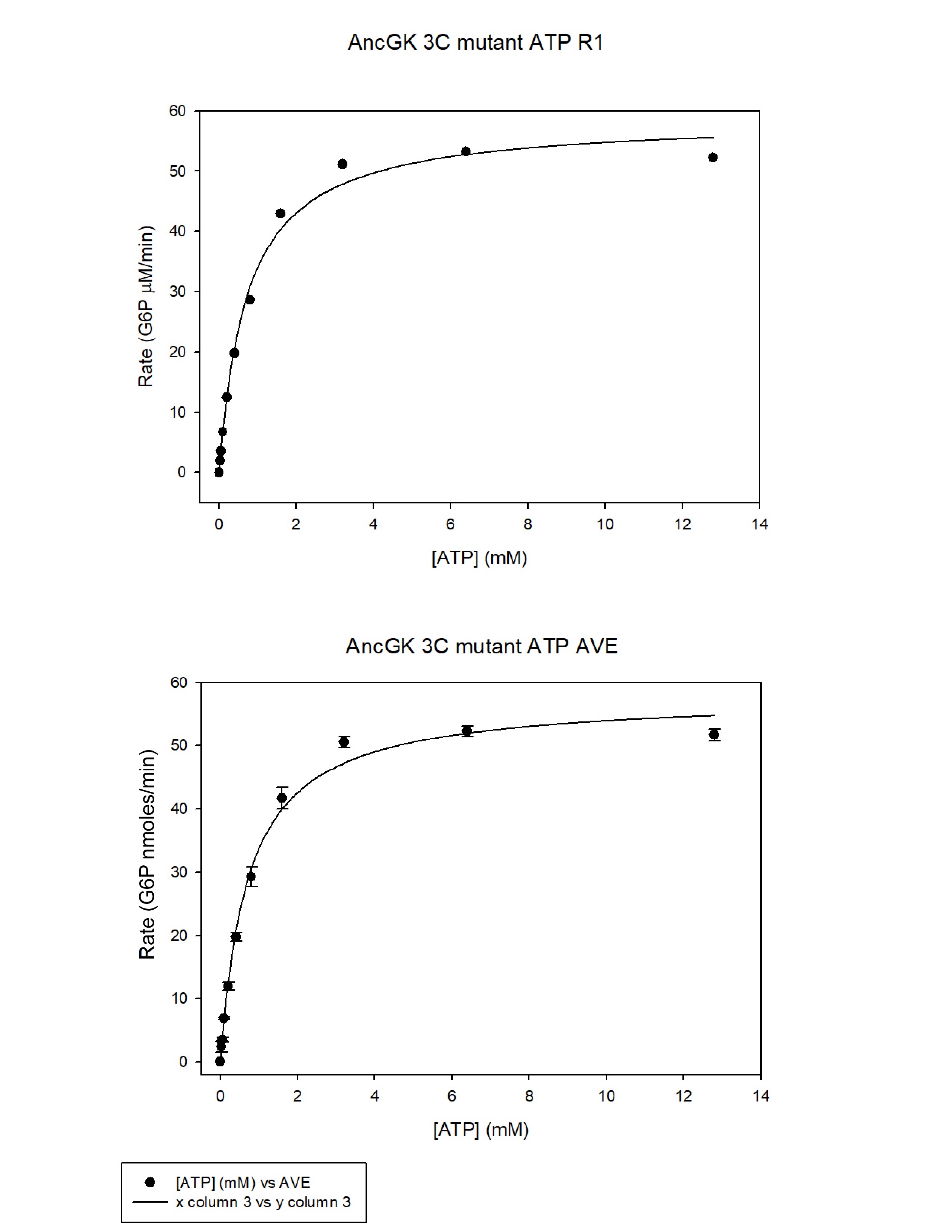

**Figure S19.** Steady-state ATP kinetic profile for C109R+C230S+ΔC461 vGCK fitted to Michaelis-Menten equation, with n = 3 technical replicates. *K_M_* = 0.71 ± 0.02 mM, *k_cat_* = 40.0 ± 0.6 s^-1^, (Ave ± SD).

**Table S1.** Primers used to construct cGCK and vGCK cysteine variants.

| **cGCK cysteine mutants** | **Primers** |
| --- | --- |
| Cysteine 109  Arginine 103 > Cysteine 103 | Forward: ‘5- gatgaagagccagatctactgtattccggaagatgttat-3’ Tm: 62.1  Reverse: ‘5- ataacatcttccggaatacagtagatctggctcttcatc-3’ Tm: 61.2 |
| Cysteine 230  Serine 224 > Cysteine 224 | Forward: ‘5- gtgggtaccggctgcaacgcgtgct-3’ Tm: 82.4  Reverse: ‘5- agcacgcgttgcagccggtacccac-3’ Tm: 82.4 |
| Cysteine 461  Inserting C at c-terminal end | Forward: ‘5- cgtctggcgtgcatgggtaagtaactcgagcac-3’ Tm: 67  Reverse: ‘5- gtgctcgagttacttacccatgcacgccagacg-3’ Tm: 67 |
| **vGCK cysteine mutants** |  |
| Cysteine 109  Cysteine 120 > Arginine 120 | Forward: ‘5-ccaaaaaccaaatgtaccgcattccg-3’ Tm: 59.2  Reverse: ‘5-cggaatgcggtacatttggtttttgg-3’ Tm: 59.2 |
| Cysteine 230  Cystine 230 > Serine 230 | Forward: ‘5- gtgggtaccggcagcaacgcgtgct-3’ Tm: 82.4  Reverse: ‘5- agcacgcgttgctgccggtacccac-3’ Tm: 82.4 |
| Cysteine 461  Deletion of cysteine at c-terminal end | Forward: ‘5- gttgcgtgcaagatggcgggtcagtaactcgagcaccac-3’ Tm: 70.3  Reverse: ‘5- gtggtgctcgagttactgacccgccatcttgcacgcaac-3’ Tm: 70.3 |
